# Learning precise segmentation of neurofibrillary tangles from rapid manual point annotations

**DOI:** 10.1101/2024.05.15.594372

**Authors:** Sina Ghandian, Liane Albarghouthi, Kiana Nava, Shivam R. Rai Sharma, Lise Minaud, Laurel Beckett, Naomi Saito, Charles DeCarli, Robert A. Rissman, Andrew F. Teich, Lee-Way Jin, Brittany N. Dugger, Michael J. Keiser

## Abstract

Accumulation of abnormal tau protein into neurofibrillary tangles (NFTs) is a pathologic hallmark of Alzheimer disease (AD). Accurate detection of NFTs in tissue samples can reveal relationships with clinical, demographic, and genetic features through deep phenotyping. However, expert manual analysis is time-consuming, subject to observer variability, and cannot handle the data amounts generated by modern imaging. We present a scalable, open-source, deep-learning approach to quantify NFT burden in digital whole slide images (WSIs) of post-mortem human brain tissue. To achieve this, we developed a method to generate detailed NFT boundaries directly from single-point-per-NFT annotations. We then trained a semantic segmentation model on 45 annotated 2400µm by 1200µm regions of interest (ROIs) selected from 15 unique temporal cortex WSIs of AD cases from three institutions (University of California (UC)-Davis, UC-San Diego, and Columbia University). Segmenting NFTs at the single-pixel level, the model achieved an area under the receiver operating characteristic of 0.832 and an F1 of 0.527 (196-fold over random) on a held-out test set of 664 NFTs from 20 ROIs (7 WSIs). We compared this to deep object detection, which achieved comparable but coarser-grained performance that was 60% faster. The segmentation and object detection models correlated well with expert semi-quantitative scores at the whole-slide level (Spearman’s rho ρ=0.654 (p=6.50e-5) and ρ=0.513 (p=3.18e-3), respectively). We openly release this multi-institution deep-learning pipeline to provide detailed NFT spatial distribution and morphology analysis capability at a scale otherwise infeasible by manual assessment.

## Introduction

Accurate identification and quantification of neuropathological hallmarks such as neurofibrillary tangles (NFTs) may be crucial for advancing our knowledge of Alzheimer disease progression and developing effective interventions[1]. Convolutional Neural Networks (CNNs) and their variants have demonstrated remarkable capabilities for image recognition and segmentation[2] tasks in the medical domain[3]. In neuropathology, deep learning on digitized whole slide images (WSIs) of brain tissue can automate detecting and quantifying distinct pathological features such as amyloid beta plaques[4–6]. This includes recognizing and quantifying the NFTs central to AD diagnosis and staging[7–10]. However, several challenges persist. Variability in staining techniques, tissue preparation, and imaging conditions across laboratories hinders the generalization of deep learning models[11,12]. Additionally, limited expert annotator bandwidth creates a scarcity of large, well-annotated datasets for neuropathologies, encouraging research in self-supervision, weak supervision, and multiple-instance learning[13–15]. Addressing these challenges is essential to deploy deep learning in neuropathology as consistent and reproducible analyses.

Building on our deep-learning-based neuropathological image analysis research[4,5,11,12,16], we introduce an open-source robust algorithm for automated detection, segmentation, and quantification of mature NFTs in the temporal lobe of AD brain tissue WSIs. Crucially, we develop a framework for automatically converting point-annotated NFTs to detailed ground-truth segmentation masks to maximize annotator bandwidth and harness active learning approaches more effectively. We leverage a carefully curated dataset from multiple Alzheimer’s Disease Research Centers (ADRCs) and employ a straightforward and reproducible modeling architecture to segment NFTs. The objective is to provide researchers with an efficient and reliable tool and framework to enhance NFT burden quantification to ultimately advance our understanding of Alzheimer disease and other neurodegenerative diseases. Quantitative data on NFT burden can aid in more robust correlations to clinical, demographic, and other data collected.

The model strongly correlates to expert-assigned WSI semi-quantitative scores on a 23-case hold-out set. These “CERAD-like” scores follow a scale for NFTs similar to that from the original CERAD criteria for neuritic plaques[5,17]. We present the methodology, including dataset curation, deep learning model architecture, evaluation metrics, and object detection benchmarks. Finally, we discuss the approach’s clinical implications and potential benefits in the broader neuropathology research context of harnessing computational methods for more accurate and consistent analysis.

## Methods

### Dataset Curation

#### Cohort selection

We obtained de-identified autopsy brain tissue samples devoid of personal identifiers and compliant with HIPAA regulations consistent with previous practices[4,5,11,12,16,18]. The annotated dataset comprises a subset of 23 cases from 295 cases collected from three distinct Alzheimer’s Disease Research Centers (ADRCs): the University of California Davis ADRC, the Columbia ADRC, and the University of California San Diego ADRC, following published case-specific inclusion/exclusion criteria[18]. These cases came from a diverse pool of research subjects recruited from various sources, including the practices of participating neurologists and community-based recruitment. The source publication delineates additional recruitment strategy information[18]. All cases met the pathological criteria for Alzheimer disease (AD), meeting NIA Reagan or NIA-AA intermediate/high criteria[19,20]. Ethical approval for this study was granted by the Institutional Review Boards at the home institutions of each ADRC, and written consent was obtained from individuals both during their lifetimes and posthumously. As the dataset originates from ADRCs, we consistently collected select data using standardized forms from the National Alzheimer’s Coordinating Center to ensure data integrity and consistency across cases[21].

#### Histology and slide-level assessments

This study used 5-7µm formalin-fixed paraffin-embedded (FFPE) sections from the temporal cortex. These sections arose from designated anatomical regions available at each ADRC. Each ADRC prepared its FFPE sections, mounted the slides, and shipped unstained slides to the University of California Davis (UCD) for staining to minimize batch effects. As previously published[18], we performed all antibody staining procedures under laboratory best practice standards, meeting Federal, State of California, and UC Davis guidelines and regulations. We used appropriate positive and negative controls for each antibody in each run.

We stained temporal cortex slides with the AT8 antibody (1:1000, Thermo Scientific, Waltham, MA, USA). All slide sections were digitized, capturing whole slide images (WSI) using a Zeiss Axio Scan Z.1 microscope at 40x magnification, creating images with a 0.11 µm/pixel resolution saved in the proprietary Carl Zeiss (.CZI) format.

An expert (BND) performed semi-quantitative histopathological assessments of NFTs on each WSI, blinded to demographic, clinical, and genetic information. The assessments followed semi-quantitative protocols outlined by the Consortium to Establish a Registry for Alzheimer’s Disease (CERAD)[17]. This protocol consisted of denoting the densest 1mm^2^ area of NFTs per WSI as none (no NFTs present), sparse (0-5 NFTs), moderate (6-20 NFTs), or frequent (greater than 20 NFTs).

### Data annotation (annotations version one)

We used Zen Blue 3.2 software for WSI annotation. We visually explored the tissue and identified three regions of interest (ROIs), each measuring 10,680 x 21,236 pixels with a resolution of 0.011 microns per pixel. We designated two gray matter ROIs that spanned the entire cortical region and one ROI along the gray matter-white matter junction (e.g., Fig. 1a). The annotation process referred to the visualizations and descriptors outlined in Moloney et al. 2021[1] to determine whether a neuron met the criteria for being a neurofibrillary tangle. While Moloney et al. 2021 define criteria for identifying pre-tangles, mature tangles, and ghost tangles, we focused solely on marking mature tangles within a cell with a clearly defined nucleolus. The trained annotator (KN) meticulously scanned the ROI, looking for flame-shaped tangles that conformed to the shape of the respective neuron defined by a visible nucleolus.

**Figure 1.**
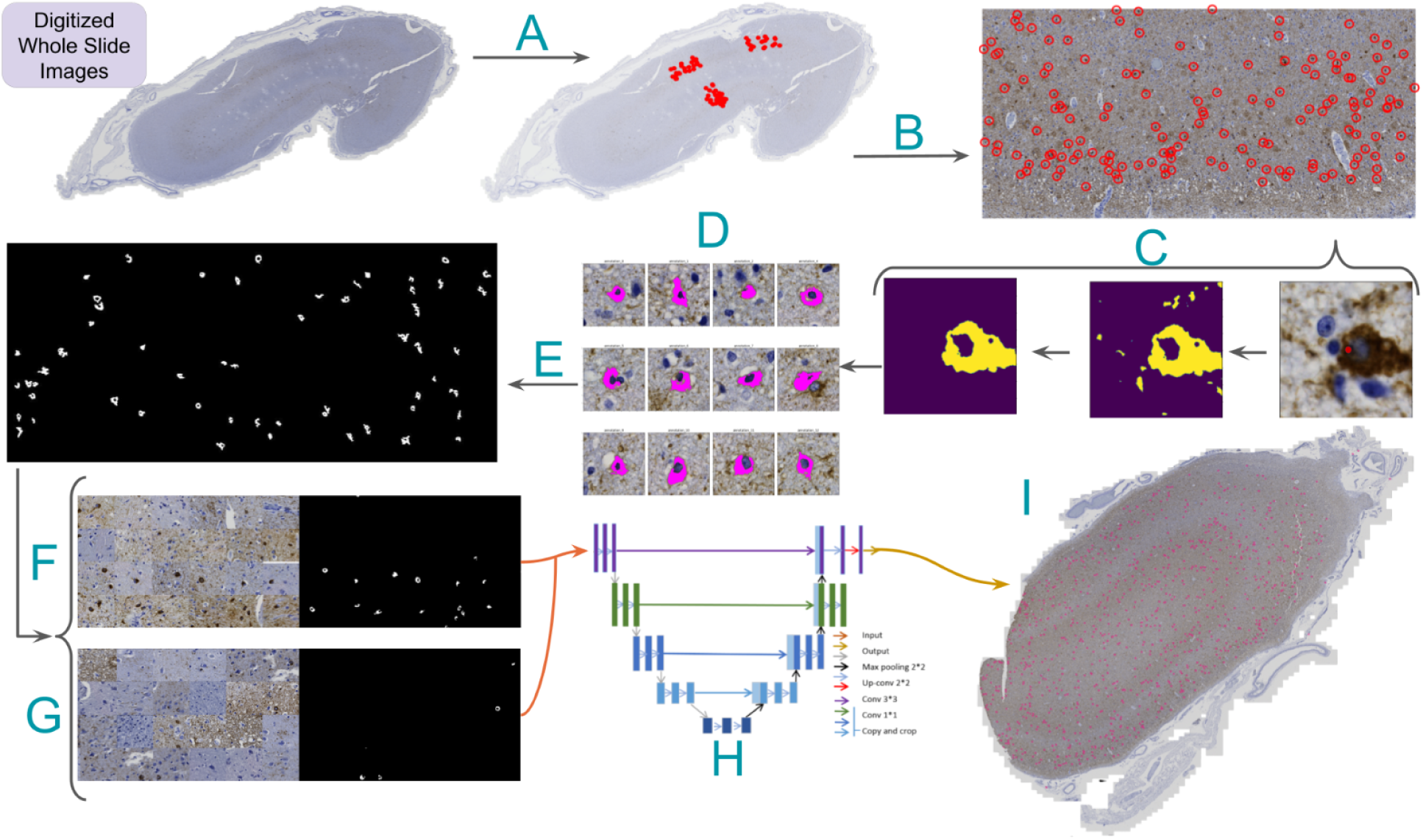
Neurofibrillary tangle model annotation and training pipeline. A) Representative slide with three regions of interest and their point-annotated NFTs. Processing steps are as follows: B) Isolate rotated ROIs and transform global NFT coordinates to local coordinates. C) Zoom to point annotation (right), isolate the DAB channel (middle), and apply morphological operations and Otsu thresholding to segment large “blobs” (left). D) Apply a center bias to remove off-center NFT candidates, retaining a single blob per tile. E) Stitch tiles into a single large ground truth mask for each ROI, replacing point annotations. F) Generate class-balanced batches for training via random NFT and tile sampling from the ROI using an input tile size of 1024x1024 pixels. G) Slice WSIs in a structured format with a stride equal to tile size. H) Feed tiles into UNet with ResNet50 backbone to generate prediction masks. I) Stitch tile predictions and overlay them onto the WSI to visualize the heatmap.

We annotated neurofibrillary tangles that exhibited defined boundaries with smooth curves, contained a nucleolus, had AT8 staining filling the cell, were located in gray matter, and typically featured 1-2 protrusions. The NFT was marked at the nucleolus (Fig. 1b) if and only if it met all criteria; otherwise, it was left unmarked. Identifying the nucleolus with a cross marking ensured a consistent size and relative location, which was crucial for subsequent training. We used a clear-cut framework following a uniform NFT definition to enhance consistency in the automated detection model (Supp. Fig. 1). A total of 1476 NFTs were annotated across the 74 ROIs.

### Datatype conversion

We converted the Carl Zeiss proprietary format images into the open-source Zarr file format to facilitate data analysis pipelines[22]. We loaded the highest resolution view of each WSI into an intermediary numpy[23] array in memory and then saved it to disk in Zarr. We set the storage chunk size to 5000 pixels in X and Y dimensions. When images had multiple scenes, we separated them into distinct files with the same base name and a suffix indicating the corresponding scene (e.g., 1-343-Temporal_AT8_s1). The Carl Zeiss format’s native compression method (JPEG XR) saves disk space at the cost of read and write speed. We used a lossless compressor (Blosc-zstd, clevel=5, bit shuffling enabled) that achieves significantly higher read and write speeds during WSI-level segmentation at the cost of disk space usage. Other users can change this compressor to suit their needs. We provide WSIs in the data repository in Zarr using the JPEG XR compression algorithm to reduce disk space and transfer bandwidth. Users will benefit from recompressing the files with the Blosc-zstd compressor for faster processing.

### Rotated ROI correction

Annotated region of interest (ROI) orientations within WSIs were not guaranteed to align with slide or image edges. This introduced complexity when loading the ROI, as arrays require slicing along fixed columns and rows parallel to boundaries. Ensuring the isolation of the ROI was crucial to prevent the inadvertent introduction of non-annotated NFTs into the dataset while preserving the total count of manually annotated NFTs. We first sliced the minimum inscribing region around the rotated ROI to address this. Second, we used the skimage library’s transform.warp method to crop the ROI. We stored each corrected ROI as a Zarr file and attached its path to a custom WSIAnnotation Python object, which was saved to disk using the pickle format for easy downstream processing[24].

### Point annotation to segmentation masks

We procedurally converted NFT point annotations to ground truth NFT pixel-boundary masks to train a semantic segmentation model. Bootstrapping point annotations into masks saves expert annotators time and scales to more annotations, facilitating efficient expert pathologist active-learning iterations. Additionally, this approach unlocks retrospective analysis for old datasets, reducing the annotation starting requirement from bounding boxes or masks to simple point annotations. Semantic segmentation enables detailed morphological analyses, WSI counts, and spatial distribution analyses of NFTs[10].

We began this procedure by cropping 400x400px tiles around each NFT in the dataset (Fig. 1c, Fig. 2a). We padded NFTs at the ROI boundary to ensure consistent centering of the NFT in the cropped tile. Each tile, centered on an NFT, underwent a custom conventional image-processing segmentation pipeline to generate a 400x400px binary mask. This pipeline involved color deconvolution using skimage’s color.rgb2hed method to convert the tile to HED color channels, followed by min-max normalization for enhanced diaminobenzidine (DAB) channel extraction[25]. The tile was then Otsu thresholded and binarized, followed by post-processing with morphological opening and closing operations[26] (Fig. 2b), isolating the AT8 immunohistochemically (IHC) stained tissue.

**Figure 2.**
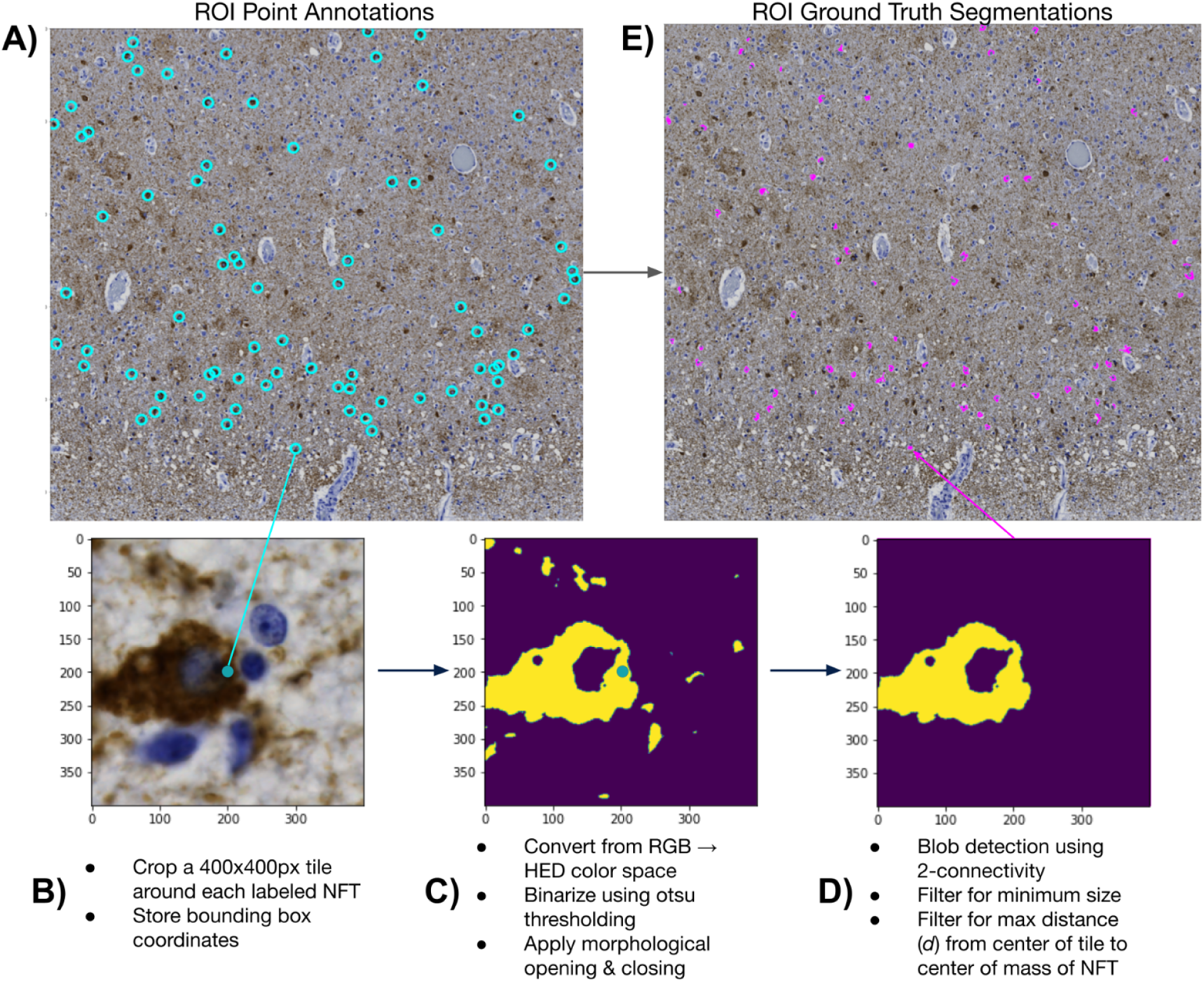
NFT point-to-mask pipeline detail. A) We convert ROIs with human NFT point annotations to detailed segmentation masks. We do so by B) generating 400x400 pixel tiles centered on the label, C) applying color deconvolution, otsu thresholding, and morphological cleaning operations, and D) performing blob detection with center-biasing and size filtering to obtain a mask. E) We then stitch each mask into its corresponding location in the ROI via union operations.

Certain tiles posed challenges, such as those having closely clustered NFTs or background noise due to the high density of phosphorylated tau protein in the surrounding tissue. To specifically segment the target NFT, we used skimage’s measure.label method to label contiguous regions of the IHC mask. Subsequently, we iteratively removed all labeled regions (blobs) whose center of mass was not within 80px of the tile’s center. This “center bias” segmentation procedure favored the NFT most centrally located within the tile. Finally, we identified the largest blob in this region, and we removed all other blobs <50% of its pixel size (Fig. 2c). We plotted the distribution of ground-truth NFT segmentations across all ROIs to check for outlying tiles and verified no tiles had mask sizes much smaller than expected (close to 0) (Supp. Fig. 2). Finally, we employed union operations and the coordinates of each tile to stitch together all masks corresponding to a single ROI upon an empty numpy array with a shape equal to its corresponding ROI, creating an ROI mask with each NFT segmented rather than point-annotated (Fig. 1e, 2d).

We applied this procedure to each WSI in the dataset (n_train/test_ = 22, n_val-batch_ = 26). We encapsulated all relevant annotation information for a WSI into a WSIAnnotation object and then saved it to disk using Python’s native object serialization library, pickle[27]. We generated lightweight pickle objects by employing Zarr for both the cropped and ground truth ROIs — we stored independent Zarr arrays and simply referenced their paths in the pickled objects. This strategy eliminated the requirement for time-consuming unpickling of the WSIAnnotation during model training and evaluation. While we successfully parallelized the creation of WSIAnnotation objects at the WSI level, the maximum RAM capacity of a workstation can still be a bottleneck. Each worker loaded a distinct WSI; considering each worker’s large memory footprint, this constraint was a significant factor.

### Model training and evaluation

We split the dataset 80/20 into training (n=15) and hold-out test (n=7) datasets stratified by WSI. We further split the training dataset into static training (n=10) and validation (n=5) sets in the same proportions stratified by WSI (Supp. Table 8).

To load tiles and their segmentation masks from disk efficiently, we designed a custom PyTorch[28] Dataset combining Zarr with weighted random sampling. We created a static data frame with rows containing coordinates generated with a stride of 1024 (equivalent to the tile size) and mappings to a single ROI/WSI pair. We assigned it a positive binary label if an NFT point annotation was present within a 1024x1024 tile loaded at the given coordinate. We employed a custom PyTorch weighted random sampler with 50/50 class balancing during training. These oversampled tiles contain NFTs across all ROIs in the training set, addressing the sparsity of NFT tiles when randomly sampled. For instance, with a batch size of 32 in our training dataloader, we generated 32 1024x1024px tiles from random locations across all training set ROIs, ensuring that, on average, 50% contained NFTs instead of including many empty ground truth tiles. In validation, testing, and inference, we loaded tiles across ROIs non-randomly to guarantee full coverage of the input image.

We deployed Kornia’s geometric and color augmentation procedures to increase our training set size and enhance the model’s robustness on the hold-out test set[29]. Kornia utilizes GPU processing to generate these augmentations and dynamically considers which transformations should apply to the image and mask, respectively (e.g., masks should not be ColorJittered). Augmentations included horizontal and vertical flips, affine transformations (e.g., rotation, rescaling, shear), color jiggle of saturation, hue, brightness, and contrast, and Gaussian blurring.

We chose a standard UNet model with a ResNet50 encoder, pre-trained on ImageNet[30–32]. We ImageNet normalized the input images to account for this pretraining. Due to the imbalanced nature of the ground truth masks (i.e., NFTs take up much less than 50% of a tile), we desired a loss function that allowed for weighing false negative classifications more heavily. We chose Tversky loss with alpha and beta parameters defined by hyperparameter optimization via Weights and Biases’ (W&B) sweeps[33,34]. We monitored the training and validation losses, F1 scores, and positive IOUs using the W&B integration with Pytorch Lightning[35].

We conducted training across two NVIDIA RTX 3090 GPUs using Pytorch Lightning’s distributed data-parallel scheme. We selected hyperparameters using W&B’s Bayesian optimizer and Hyperband early-stopping procedures[36]. Sweep parameters included learning rate, weight decay, optimizers, momentum, and alpha parameters for the Tversky loss function (Supp. Fig. 3a). We tested on a single GPU with non-overlapping tiles to avoid potentially duplicated samples. Key metrics for assessing the best model performance included positive IOU, F1 score, and mIOU. We chose the epoch with the lowest validation loss for the final evaluation of the test set.

### Agreement maps

We visualized model performance across varying fields of view to better understand the model’s selection criteria for NFTs. We thresholded predictions at 0.5 to convert pixel values from continuous to binary class labels that correspond to whether an NFT is present (value of 1) or absent (value of 0). We then constructed ‘agreement maps’ by directly comparing pixel-wise values between the ground truth and predictions (Fig. 3). Pixels are ‘true positive’ (TP) if their ground truth and predicted value at a given index are equal to 1 and ‘true negative’ (TN) if the ground truth and predicted values are 0. We assigned ‘false positive’ (FP) labels where the model incorrectly predicts an NFT that is not present in the ground truth (i.e., ground truth pixel value = 0, predicted pixel value = 1) and ‘false negative’ (FN) labels to the inverse. We used Matplotlib’s LinearSegmentedColormap class to distinguish TP, FP, and FN masks[37]. All other regions of the ROI, i.e., those without annotated NFTs, were considered TN.

**Figure 3.**
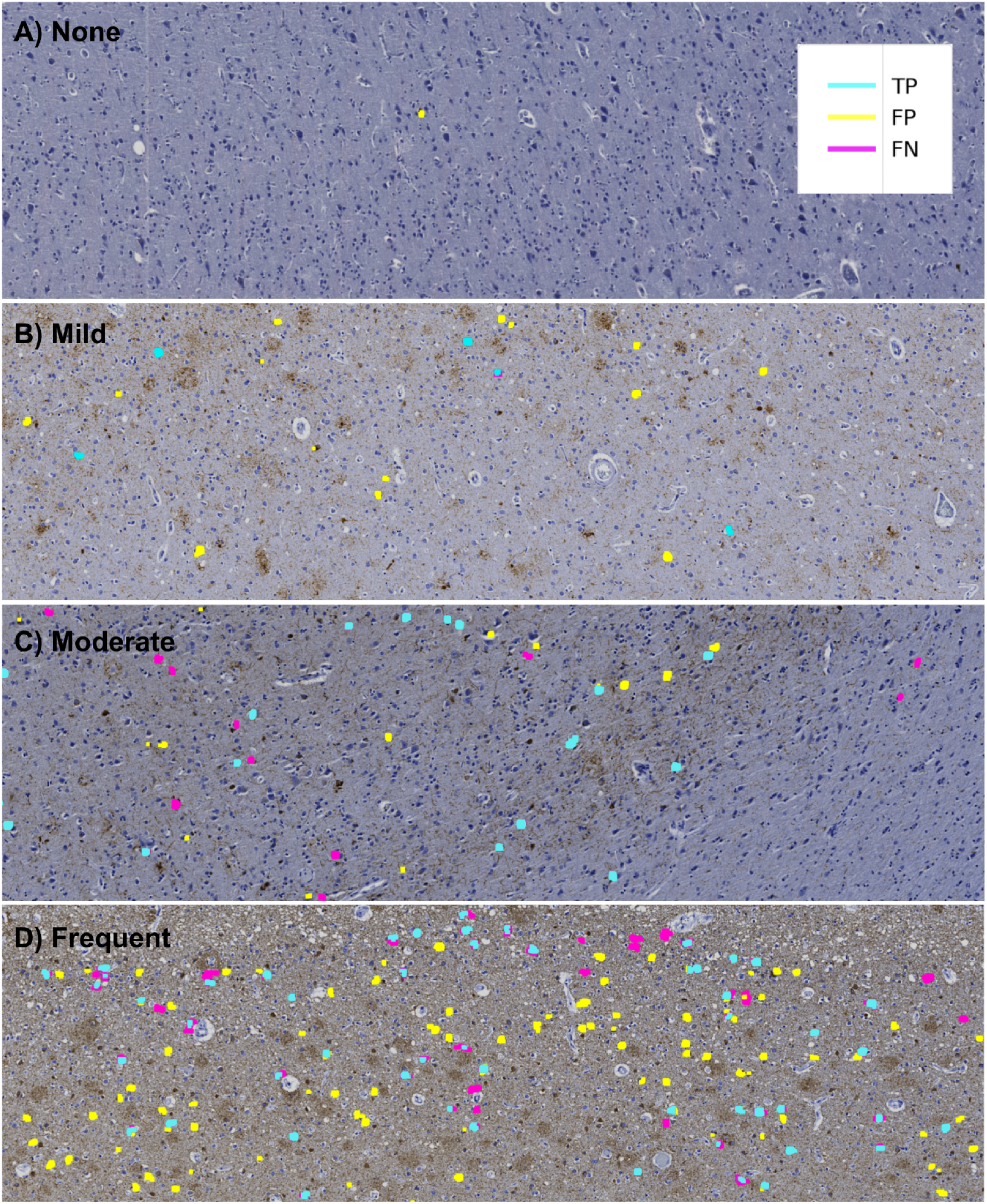
Agreement maps display how NFT predictions compare to human labels. Agreement maps from four different ROIs in order of increasing semi-quantitative severity. As the slide-level severity score increases, the model detects more NFTs. However, False Positives (FPs, yellow) and False Negatives (FNs, magenta) increasingly appear as the severity increases. (A) None, (B) Mild, (C) Moderate, and (D) Severe CERAD-like NFT burden scoring by expert assessment. Notably, True Positive (TP, cyan) predictions display high pixel-wise boundary fidelity, and disagreement between human labels and model predictions more typically applies to whether the entire object qualifies as a mature NFT.

### Data re-annotation (annotations version two)

When examining preliminary results from the agreement maps, we observed many FP masks across the ROIs throughout the entire dataset (Fig. 3c, 4a). A closer examination of the FP-labeled raw image areas revealed many plausible predicted NFTs meeting the annotation criteria described earlier (which was carried out by a trained novice) (Fig. 4a, 4c). To stress-test the correctness and validity of our ground truth data, we initiated a re-annotation experiment conducted by an expert (BND) to rescue any NFTs overlooked in the initial round of annotations.

**Figure 4.**
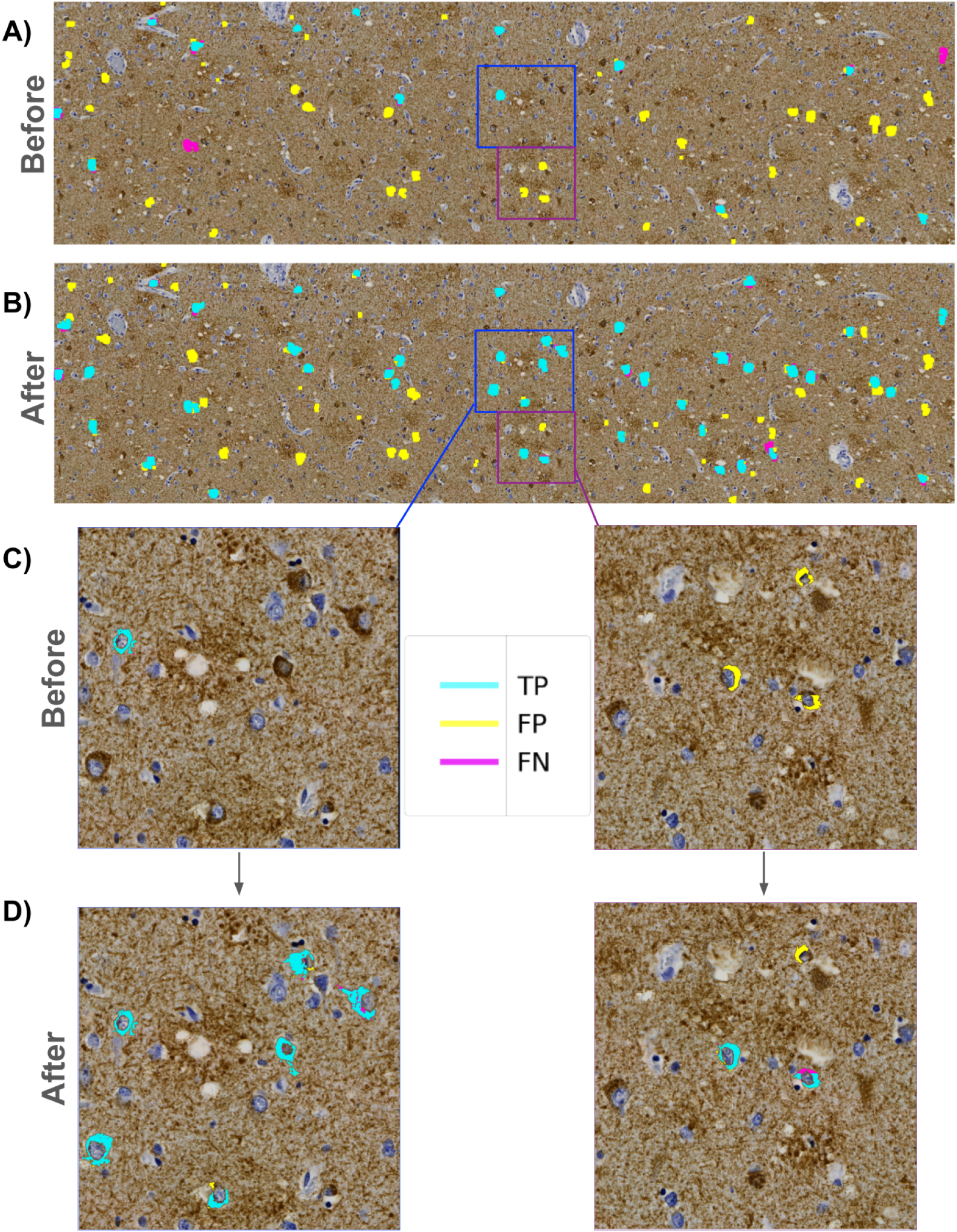
Expert reannotation rescues putative predicted NFTs. (A) ROI-level agreement map comparing trainee-annotated ground truth NFT labels versus “version 1” model-predicted segmentation masks. (B) Agreement map of the same ROI using expert-reannotated ground truth NFT labels versus “version 2” model-predicted masks. C) Zoomed view highlighting initial trainee-vs-model agreement maps. D) Zoomed view highlighting annotated expert-vs-model agreement maps. Multiple newly expert-annotated (“rescued”) NFTs that were previously labeled FPs or TNs become TPs after integrating the reannotated data and retraining the model.

We sliced each ROI into fifteen 4247x3560 pixel “super-tiles,” which were large enough for our expert annotator to process efficiently while maintaining sufficient resolution to evaluate NFTs displayed in the super-tiles. We uploaded all super-tiles to the SuperAnnotate[38] platform in a randomized order. Each re-annotation study image consisted of two identical side-by-side super-tiles — with the raw super-tile image on the right and the same image overlaid with the agreement map labels on the left (Supp. Fig. 4). We chose this approach to accommodate annotator bandwidth limitations. As the focus was on correcting potentially mislabeled FP objects, the expert annotator concentrated on objects with FP masks, using SuperAnnotate’s point annotation tool to rapidly annotate any object meeting the NFT annotation criteria described earlier. No objects/NFTs were removed during the re-annotation experiment.

Following the expert’s examination of the super-tiles in the re-annotation procedure, we downloaded the re-annotation data from SuperAnnotate, which included JSON files for each image with new point annotations. We parsed these files and generated a table containing the WSI and ROI IDs, super-tile coordinates relative to the ROIs they originated from, and the coordinates of the new point annotations. In total, 280 new point annotations (a 19% addition overall) across 17 WSIs were added in the re-annotation experiment, providing higher-quality ground truth labels for training and evaluating the model (Fig. 4b, 4d). The entire dataset was re-annotated by BND in less than 90 minutes, facilitated by the platform and the outlined scheme.

### Correlating WSI-level model scores with semi-quantitative scoring

We performed rapid inference using structured slices (sliding window approach) to construct WSI-level segmentations with the model. This allows for manual inspection of segmentations across the entire WSI by overlapping the produced segmentation mask and the original WSI (Fig. 5a). Subsequently, we generated a score summarizing the image’s NFT burden. We opted for a count-based score, which tallies the total number of contiguous blobs in a downscaled image using skimage’s measure.label function. The downscaled tissue area then normalizes this count to ensure consistent burden scores. The tissue area is calculated by applying histomicsTK’s[39] tissue detection algorithm to isolate a binary mask and then summing the pixel area. Finally, we rescaled scores by the median tissue area of the training set to produce an interpretable result. We refer to this score as an NFTDetector score.

**Figure 5.**
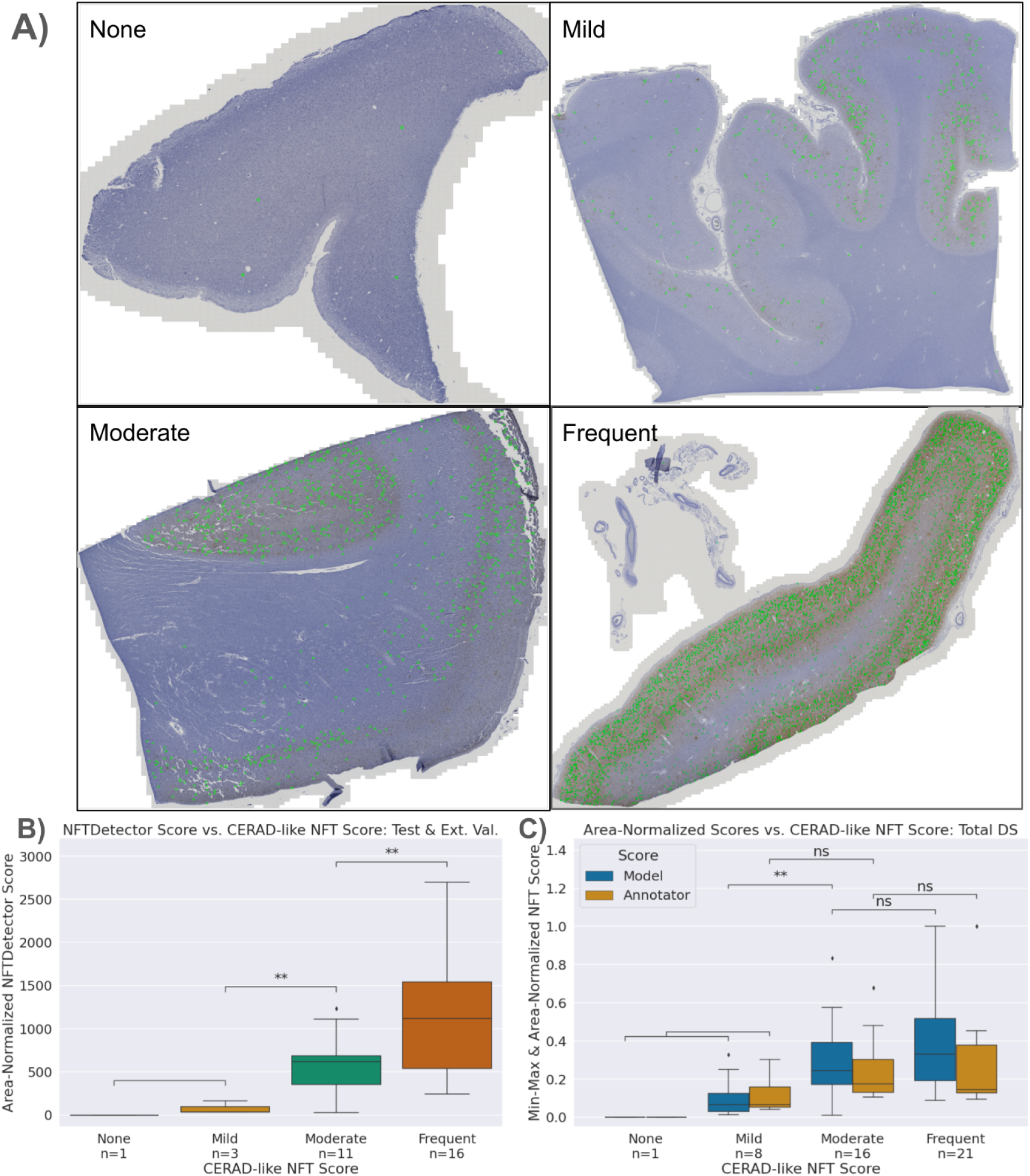
Model-derived NFTDetector scores correlated with expert CERAD-like scores. A) Representative WSIs spanning the four semi-quantitative stages of increasing NFT burden severity, ordered by CERAD-like scoring. Model NFT detections overlaid in green. B) Area-normalized NFTDetector scores on test set WSIs correlate with semi-quantitative ordinal categories. C) Area-normalized NFTDetector scores calculated on the full WSI dataset versus area-normalized counts for the same WSIs derived from the annotator labels. **: 0.001 < p <= 0.01 by Welch’s t-test. We used different assessment datasets in B) vs C) because the external test set lacks human NFT point annotations.

We ran all slides in the training, validation, and testing datasets and a separate hold-out batch of images (not point-annotated, only having slide-level scores) through the model to generate a distribution of NFTDetector scores (Supp. Table 7, 9). We then assign the WSIs and their NFTDetector scores into groups corresponding to their slide-level CERAD-like scores assigned by an expert pathologist (four grades ranging from None to Severe) and tested for statistical difference via Welch’s t-test using the statannotations^3^ library (Fig. 5b).

We compared the model’s slide-level semi-quantitative correlative performance with the trainee’s by applying the same score generation logic directly to the ground truth point annotations. By counting the number of point annotations across all ROIs for a given WSI and then normalizing by the total pixel area of the ROI, we generated a human analog to the NFTDetector score. We scaled scores by the median tissue area and min-max normalized to achieve better alignment of the y-axis for plotting purposes only. The null hypotheses aimed to validate whether the trainee’s semi-quantitative score groups were statistically different; this was confirmed via Welch’s t-test, as scores were not guaranteed to be normally distributed (Fig. 3b).

### Comparing model predictions with AT8 burden and semi-quantitative scores

We generated simple AT8 burden scores by calculating the proportion of DAB signal present in an ROI. To do this, we applied a similar pipeline to the initial steps of the point-to-mask pipeline. We first converted the image from RGB to HED channels via skimage, then binarized the image using Otsu thresholding (Supp. Fig. 5a). We then counted the total number of detected DAB positive pixels and normalized by the total number of pixels in the grayscale image to generate a DAB Proportion score (Supp. Fig. 5b). We used this DAB Proportion score and the expert-assigned semi-quantitative CERAD-like scores as correlative variables with the models’ performance metrics (Supp. Fig. 5b, 5c). We constructed boxplots and scatterplots with seaborn and tested for significant differences with statannotations[40,41].

### Statistical methods

This study operates at any of four levels. From the most to least granular: individual NFT objects (n_test_=664), tiles (n_test_=4620), ROIs (n_test_=20), and WSI images (or decedents, n_test_=7). Only the object detection analyses (YOLOv8, segmentation bounding boxes) can operate at the level of individual NFTs (“object-level”). We use this level for Figure 7 or whenever explicitly denoted. We use a tile-level loss function to train the segmentation and object detection models, reported in Supp. Fig. 3. We report ROI-level metrics in Figure 6b because the ground truth only exists at the ROI level. Performance and other characteristics are available in detail at the ROI level (Supp. Tables 3-6). Finally, at the WSI level (1-to-1 with decedent), we report correlations against semi-quantitative scores as in Figures 5b, 5c, and 7c (Supp. Table 7). The sample sizes for the training data were as follows: objects (n_train_=792), tiles (n_train_=13860), ROIs (n_train_=45), and WSI images (or decedents, n_train_=15).

**Figure 6.**
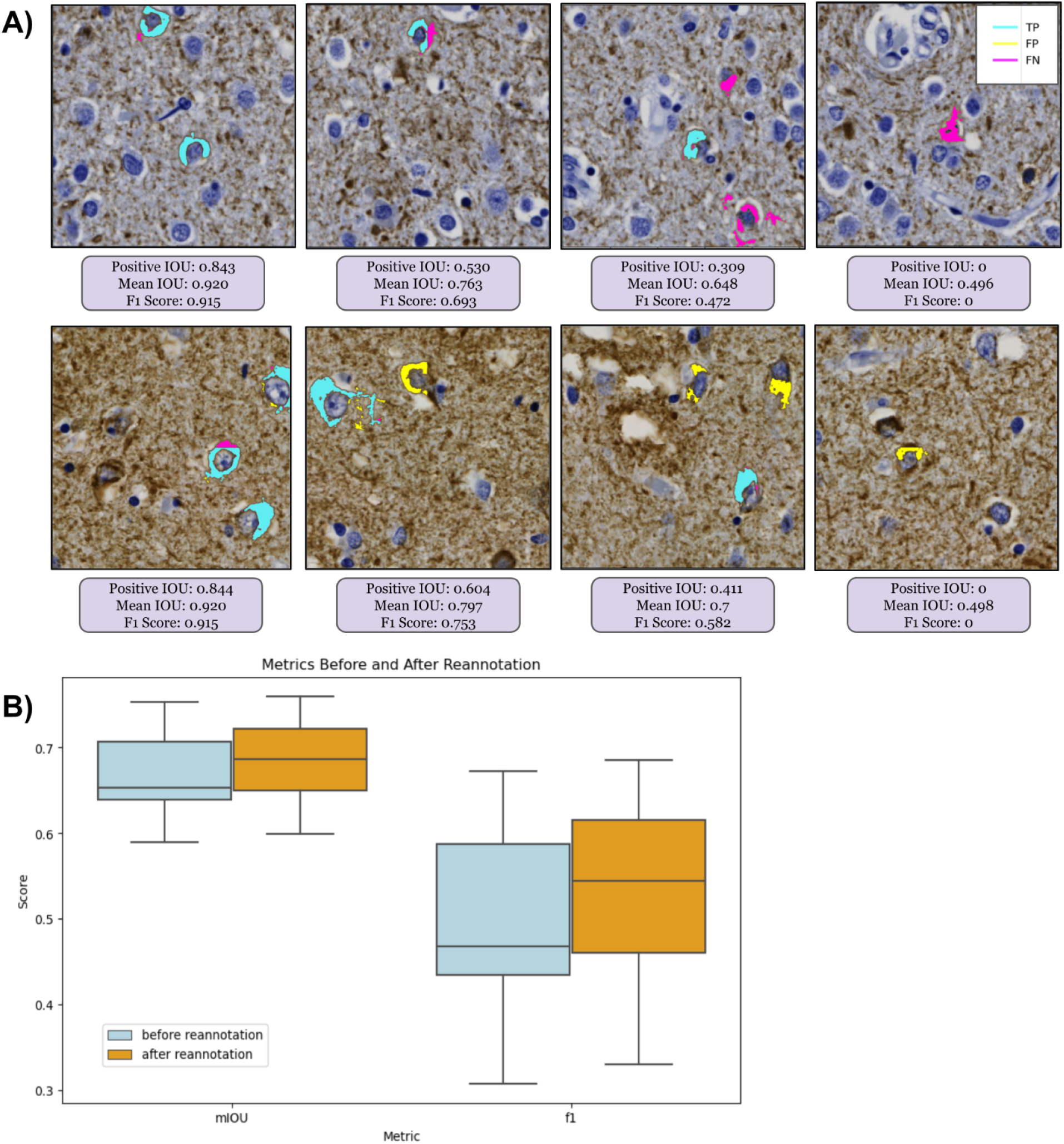
Segmentation numerical performance metrics were not always visually intuitive. (A) Progressing in order of decreasing performance (left to right): Four 1024x1024 pixel tile-level agreement maps with corresponding numerical performance metrics for segmentation performance at the NFT object level from two different ROIs (rows). The F1, mean, and positive IOU scores are specific to each 1024x1024 tile. IOU and F1 scores suffer the most from false negatives, even when the model predicts other NFT pixels correctly. (B) At the ROI level, model performance on the test set, assessed by F1 and mean IOU, improved after reannotation, with substantial qualitative improvement evident by visualization.

**Figure 7.**
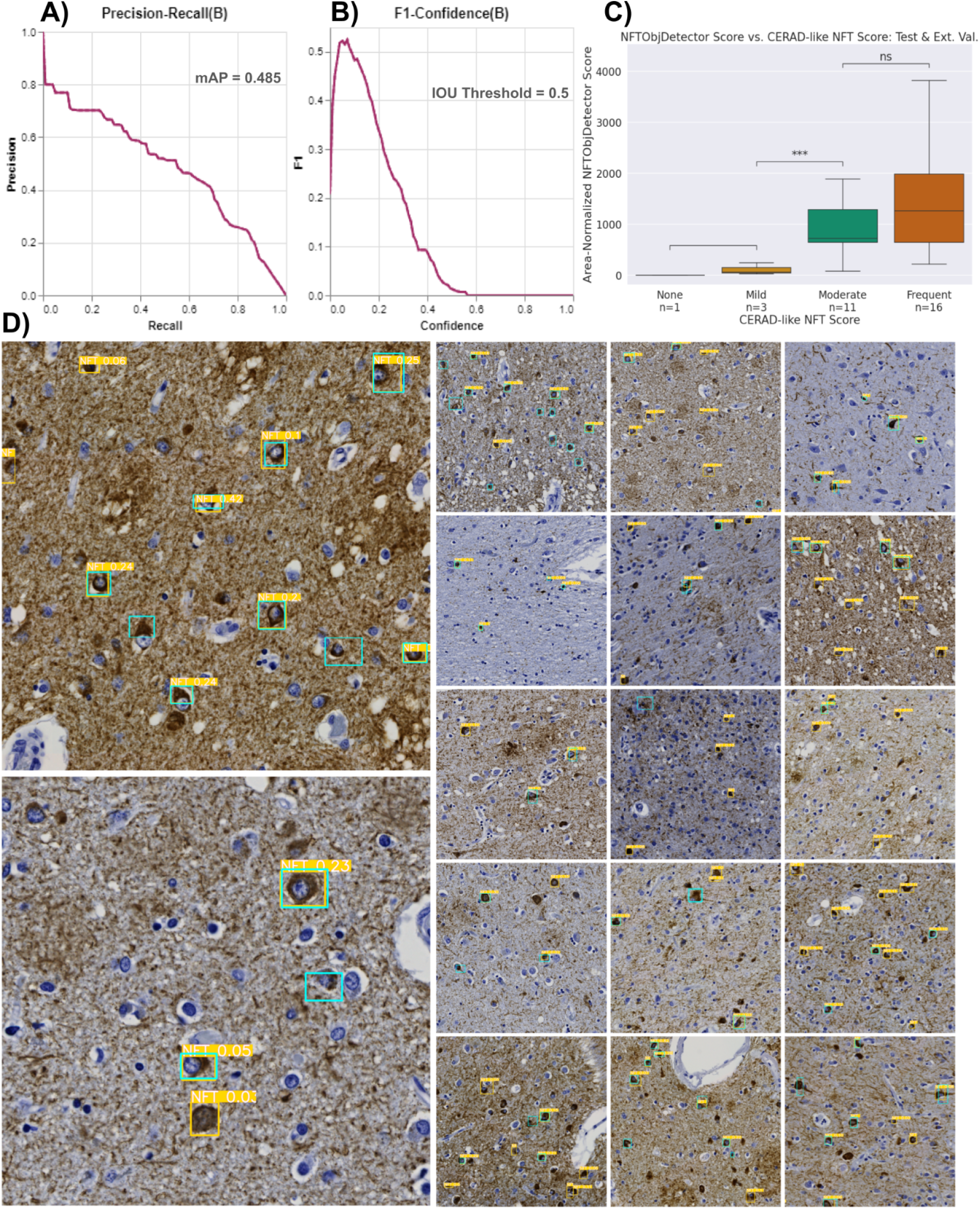
Repurposing the NFT dataset for object-detection models yields similar overall performance to the segmentation approach. A) The precision-recall curve plots the tradeoff between the YOLO model’s object-detection precision and recall across all potential confidence thresholds aggregated across all objects in the test dataset. B) Plot of aggregated object-wise F1 versus confidence threshold; low (permissive) thresholds yielded the highest F1 scores. C) WSI-level NFT counts from the YOLO model correlated with CERAD-like semi-quantitative scoring, similar to the segmentation-derived NFTDetector scores in Figure 5b. **: 0.001 < p <= 0.01 by Welch’s t-test. D) Example tiles from different ROIs overlaid with ground truth (cyan) and predicted (yellow) bounding boxes. The values highlighted in yellow are confidence scores for the predictions. Reference B) to determine which NFTs would be detected.

To calculate statistical significance, we proceeded as follows. In Table 1, we tested continuous and normally distributed variables using one-way ANOVA, continuous and non-normal variables using Kruskal-Wallis, and categorical variables via Chi-squared (Supp. Table 2). We used Welch’s t-test to compare continuous vs. categorical variables, as in Figures 5b, 7c, and Supp. Figs. 5b-c, and 9, and student’s t-test for Supp. Fig. 10a. Combining annotated (n=21) and model (n=44) scores of 44 decedents, we assessed score differences among Severe, Moderate, and Mild CERAD-like categories and score type (annotator vs. model). We excluded the None CERAD-like category due to insufficient examples (n=1); however, both the annotator and the model correctly assigned this slide. To meet model assumptions, we used a mixed-effect model and transformed scores using the natural logarithm. For Figure 6b, we use linear mixed models (Statsmodels.formula.api mixedlm [42]) to estimate the mean values and 95% confidence intervals for ROI-level metrics (e.g., pIOU, F1, and mIOU) from 7 decedents in the test set with repeated samples from each decedent, typically corresponding to 3 ROIs.

**Table 1:** Demographics grouped by dataset split.

|  | Grouped by Dataset Split |  |  |  |  |  |  |
| --- | --- | --- | --- | --- | --- | --- | --- |
|  |  | Missing | Overall | Test Set | Training Set | Validation Batch | P-Value |
| n |  |  | 45 | 7 | 15 | 23 |  |
| Sex, n (%) | Female | 0 | 26 (57.8) | 3 (42.9) | 9 (60.0) | 14 (60.9) | 0.684 |
|  | Male |  | 19 (42.2) | 4 (57.1) | 6 (40.0) | 9 (39.1) |  |
| Age at Death, Yrs median [Q1,Q3] |  | 0 | 83.0 [77.0,88.0] | 83.0 [78.0,86.0] | 84.0 [78.0,90.0] | 82.0 [78.0,86.5] | 0.574 |
| Hispanic ethnicity, n (%) | Yes | 0 | 13 (28.9) | 1 (14.3) | 4 (26.7) | 8 (34.8) | 0.562 |
|  | No |  | 32 (71.1) | 6 (85.7) | 11 (73.3) | 15 (65.2) |  |
| Hispanic origins, n (%) | Cuban | 32 | 1 (7.7) |  |  | 1 (12.5) | 0.785 |
|  | Dominican |  | 2 (15.4) |  | 1 (25.0) | 1 (12.5) |  |
|  | Mexican |  | 4 (30.8) |  | 1 (25.0) | 3 (37.5) |  |
|  | Puerto Rican |  | 3 (23.1) |  | 1 (25.0) | 2 (25.0) |  |
|  | Unknown |  | 3 (23.1) | 1 (100.0) | 1 (25.0) | 1 (12.5) |  |
| Race, n (%) | African American | 0 | 1 (2.2) |  | 1 (6.7) |  | 0.496 |
|  | Other |  | 4 (8.9) |  | 2 (13.3) | 2 (8.7) |  |
|  | Unknown |  | 3 (6.7) |  | 2 (13.3) | 1 (4.3) |  |
|  | White |  | 37 (82.2) | 7 (100.0) | 10 (66.7) | 20 (87.0) |  |
| Years of Education, median [Q1,Q3] |  | 0 | 15.0 [12.0,16.0] | 14.0 [12.5,15.5] | 16.0 [10.5,16.0] | 14.0 [12.0,16.0] | 0.902 |
| CERAD-like* NFT Score, n (%) | 0 | 0 | 1 (2.2) | 1 (14.3) |  |  | 0.044 |
|  | 1 |  | 8 (17.8) | 1 (14.3) | 5 (33.3) | 2 (8.7) |  |
|  | 2 |  | 16 (35.6) | 4 (57.1) | 5 (33.3) | 7 (30.4) |  |
|  | 3 |  | 20 (44.4) | 1 (14.3) | 5 (33.3) | 14 (60.9) |  |
| PMI, median [Q1,Q3] |  | 34 | 6.0 [2.8,21.9] | 24.0 [22.4,36.0] | 6.0 [3.1,14.5] | 5.2 [0.0,6.0] | 0.057 |
| Primary Clinical Diagnosis, n (%) | AD | 2 | 20 (46.5) | 4 (57.1) | 4 (30.8) | 12 (52.2) | 0.169 |
|  | CERAD Definite AD |  | 2 (4.7) | 1 (14.3) | 1 (7.7) |  |  |
|  | CERAD Probable AD |  | 1 (2.3) |  | 1 (7.7) |  |  |
|  | Definite AD |  | 11 (25.6) |  | 5 (38.5) | 6 (26.1) |  |
|  | LB variant AD |  | 8 (18.6) | 1 (14.3) | 2 (15.4) | 5 (21.7) |  |
|  | Probable AD |  | 1 (2.3) | 1 (14.3) |  |  |  |
| ADRC Origin, n (%) | Columbia | 2 | 20 (46.5) | 2 (28.6) | 7 (53.8) | 11 (47.8) | 0.642 |
|  | UC Davis |  | 9 (20.9) | 1 (14.3) | 3 (23.1) | 5 (21.7) |  |
|  | UC San Diego |  | 14 (32.6) | 4 (57.1) | 3 (23.1) | 7 (30.4) |  |
| Braak NFT Stage, n (%) | III | 15 | 1 (3.3) | 1 (14.3) |  |  | 0.323 |
|  | IV |  | 4 (13.3) | 1 (14.3) | 2 (25.0) | 1 (6.7) |  |
|  | V |  | 6 (20.0) | 2 (28.6) |  | 4 (26.7) |  |
|  | VI |  | 19 (63.3) | 3 (42.9) | 6 (75.0) | 10 (66.7) |  |
| CERAD, n (%) | Sparse | 12 | 2 (6.1) |  | 1 (10.0) | 1 (5.9) | 0.496 |
|  | Moderate |  | 4 (12.1) | 2 (33.3) | 1 (10.0) | 1 (5.9) |  |
|  | Frequent |  | 20 (60.6) | 4 (66.7) | 5 (50.0) | 11 (64.7) |  |
|  | Not assessed |  | 7 (21.2) |  | 3 (30.0) | 4 (23.5) |  |
| Thal, n (%) | 4 | 31 | 1 (7.1) | 1 (25.0) |  |  | 0.559 |
|  | 5 |  | 10 (71.4) | 2 (50.0) | 3 (75.0) | 5 (83.3) |  |
|  | Not assessed |  | 3 (21.4) | 1 (25.0) | 1 (25.0) | 1 (16.7) |  |

Demographics, neuropathologic variables, and clinical diagnoses. P-values were calculated between datasets using appropriate statistical tests; continuous and normally distributed variables were tested using one-way ANOVA, continuous and non-normal variables were tested using Kruskal-Wallis, and categorical variables were tested via Chi-squared test. PMI: Post-mortem interval; LB: Lewy body.

### Metric choices

Object detection and instance segmentation models typically report mean average precision (mAP), while fewer studies report segmentation model performance by mAP. Since we perform binary segmentation, mAP is equivalent to AUPRC for the positive class[43]. We observed that the model’s confidences are highly skewed towards 0 or 1, such that varying the confidence threshold for the segmentation predictions does not meaningfully alter performance (Supp. Fig. 6). For this reason, we report F1 instead (i.e., Dice score for pixel-based calculations), as it is approximately the same as mAP given the above constraints. To determine a random F1 baseline, we used a null random model that achieves a precision equal to the test set ROIs’ positive pixel prevalence of 0.000898 (CI: [0.000205, 0.00159]) and a recall of 0.5. F1 can be further deconstructed into precision and recall at the chosen threshold, displayed within the figures of this study. We also report mean intersection over union (mIOU) — the average of positive and negative IOUs — and another standard segmentation metric. We calculate AUROC using sklearn.metrics.RocCurveDisplay [44]. We aggregate these metrics across all pixels in the dataset during training (i.e., flattening the predictions) but report the test set metrics at the ROI level.

### Object detection: Converting masks to bounding boxes for instance detection metrics

To align with Signaevsky et al. 2019[8] when comparing model performance, we transformed NFT segmentations into object detections to obtain object-level metrics, as they appear to have done in their study (we could not find precise metric details). While this assessment method may not offer the granularity of pixel-level metrics, it can be more forgiving if an entire NFT prediction does not perfectly align with its ground truth mask.

To implement this, we leveraged histomicsTK’s contour generation functions to generate bounding boxes from stitched ROI prediction masks. We merged nearby bounding boxes within 150 pixels from center to center via a custom algorithm (Supp. Fig. 7a). We generated bounding boxes around the ground truth masks by taking the minimum and maximum coordinates of the mask in a cropped 400x400px view of each point-annotated NFT, consistent with the tile size used to construct the masks (Supp. Fig. 7b). We then matched bounding boxes from the same ROI for greatest IOU. We assigned unmatched and very low IOU prediction boxes (i.e., IOU < 0.001) as false positives and unmatched ground truth boxes as false negatives. We used these values to calculate our dataset splits’ precision, recall, and F1 scores.

### Object detection: Training an object detection model from bootstrapped bounding boxes

We generated ROI-level ground truth bounding boxes as described above (i.e., min/max coordinates of ground truth masks cropped around each NFT, performed at the ROI level). We then leveraged these ROI-level bounding boxes to create tile-level bounding boxes to train a YOLOv8 model[45,46]. We assigned an ROI-level bounding box to a single tile if it contained the center of the bounding box. We saved non-overlapping 1024x1024px tiles from the ROIs as static PNG files due to a limitation of the Ultralytics library, which we used to train the YOLOv8 model[45,46]. We shifted bounding box coordinates according to their assigned tile’s coordinates, clipped them to fit within the tile, converted them to YOLO format ([label, xmin, ymin, xmax, ymax]), and then saved them into a text file with the same name as its corresponding tile PNG.

We followed the protocol to generate a YOLOv8 configuration file in YAML format. In this file, we defined the paths to the train, validation, test directories, and the names of our labels (e.g., 0: NFT). Training a YOLOv8 model was straightforward after constructing the dataset with the correct input format (see Supp. Fig. 3b for loss curves). To maximize performance, we optimized hyperparameters with the default search parameters suggested by Ultralytics and their built-in hyperparameter search evolution algorithm via model.tune() for 30 iterations. The best hyperparameter settings are included in the Github repo file, “best_hyperparameters_yolo.yaml,” and tuning results can be found in Supp. Fig. 8.

### Use of LLMs

We used ChatGPT (https://chat.openai.com/, February 2024) to provide editing feedback for the manuscript and code. We also used it to convert tables from raw text into Markdown and to generate a first draft of the Abstract from the study results and field context, which we further edited. During manuscript editing, we subjected specific sections of the text to editing review by ChatGPT to enhance clarity and conciseness, manually incorporating suggestions as appropriate.

## Results

### We automatically generated pixel-wise masks of NFTs from single-point annotations

We developed an automated point-to-mask image processing pipeline to reproducibly generate pixel-level masks (outlines) around NFTs solely from manual point annotations. The motivation was that manually outlining these objects at a sufficient scale to train a deep learning model is exceedingly time-consuming, nuanced, and prone to high degrees of interobserver variation in determining the precise boundary of an NFT. The resulting point-to-mask pipeline generated high-quality ground truth segmentations with few visually erroneous examples despite leveraging solely conventional image-processing libraries (see Methods). We sampled 10 ROIs across semi-quantitative categories to assess the generated segmentation masks’ accuracy, inspecting 50 ground truth NFT segmentations per ROI. We also empirically validated results by generating a distribution of NFT ground-truth masked pixel areas and then visually inspecting examples of the largest and smallest masks in the dataset.

Using this automated pipeline, we processed 1476 objects across the 74 ROIs. The resulting masks, on average, comprised 5.5% of a tile (or 8800 pixels in area at 0.11 microns per pixel), with a minimum of 0.25% (∼400 pixels), a maximum of 32% (51,000 pixels), and a standard deviation of 3% (4940 pixels). By this method, we proceeded through an iterative process to refine parameters of the point-to-mask pipeline, such as the center bias and minimum size variables, to ensure NFT masks were sufficiently large and to minimize the chance that they are over-segmented as sometimes seen in high-background regions (Supp. Fig. 2). In the originally calculated masks, many objects would be missed due to an over-stringent center bias parameter. We corrected this by increasing the “center bias” parameter from 80 pixels to 120 pixels. The center bias procedure ensured that the detected blob’s center of mass was within a predefined number of pixels from the center of the point annotation. This automatically fixed the ∼7% of the tiles missed in the first iteration. NFTs that were typically affected by this procedure were those with longer tails and contiguous staining from nucleus to axon. After applying the filtering pipeline, we inspected the histogram of mask sizes and found the proportion of over-segmented tiles to be acceptable (<1% of tiles). Rather than manually adjust these masks, we included them to maintain a fully automated pipeline and to determine whether the trained model would be robust to the noisier ground truth labels.

### We compared how guided expert reannotation affected segmentation model quality

As there is no single metric to score how well a model identifies (segments) an object at the pixel level, we used standard segmentation metrics of mIOU and F1. As it may not be clear from numerical scores alone what constitutes mediocre, acceptable, versus exceptional performance, we frequently inspected representative examples visually (Fig. 6a). In the “version 1” model before expert reannotation, the F1 at the ROI level was 0.49 (CI: [0.44, 0.54]). Following retraining on the re-annotated dataset, the final (“version 2”) model performance improved by about 8% at the ROI level to a precision of 0.53 (CI: [0.37, 0.7]), recall of 0.60 (CI: [0.49, 0.71]), F1 of 0.53 (CI: [0.45, 0.61]), mIOU of 0.68 (CI: [0.65, 0.72]) and AUROC of 0.832 (CI: [0.781, 0.883]) on a test set of seven held-out whole slide images (WSIs) (Fig. 6b). Additionally, the model’s area-normalized WSI score increased its correlation with semi-quantitative scores on the hold-out test sets (Spearman’s rho, ρ, increased from 0.624 to 0.654) (Supp. Fig. 9).

### Segmentation and repurposed instance-detection interpretations were consistent

Compared with other studies, we also repurposed the segmentation model predictions as though the masks only predicted objects – i.e., independently of whether the predicted NFT pixel boundaries were precisely accurate. We calculated approximate bounding boxes from the ground truth and prediction masks (see Methods) to do this. This allowed us to assess the NFT segmentation model as a coarse-grained object detector model instead. We scored NFTs as true positive (TP) if the intersection over union (IOU) between the prediction and ground truth bounding boxes exceeded 0, as reported previously in the field[47]. False positive (FP) NFTs were predictions that did not overlap with ground truth bounding boxes, while FNs were ground truth bounding boxes that did not overlap with predictions. Intriguingly, this more categorical and coarse-grained object-level performance nonetheless mirrors pixel-level performance, yielding a test set F1 of 0.53 (CI: [0.45, 0.62]; Supp. Table 1).

### A dedicated YOLOv8 model did not improve NFT instance detection

Whereas the above approach reinterpreted the NFT segmentation model *post hoc* as an NFT object detector, we posited that an established object-detection architecture trained from scratch on the bootstrapped bounding box dataset might perform better at this task. Accordingly, we trained a standalone YOLOv8 model with the sole task of object detection using the ground truth bounding boxes constructed for the object detection metric baseline. The YOLOv8 model performed comparably but no better than the repurposed segmentation model, achieving an ROI-level F1 of 0.53 (CI: [0.46, 0.60], Fig. 7b) and an aggregate object-level mAP50 (equivalent to AUPRC for the NFT class) of 0.485 (Fig. 7a). The model also displayed similar effectiveness in categorizing slides into “CERAD-like” NFT semi-quantitative grades (ρ = 0.513, Fig. 7c). Visual inspection revealed that FPs were predominantly NFT-like objects failing to meet our particular annotation criteria. At the same time, FNs likely resulted from this specificity (Fig. 7d).

### Performance did not vary by AT8 stain burden or CERAD-like category

We noticed that the point-to-mask pipeline generated larger masks from dark AT8 staining around cropped NFTs, so we posited that some ground truth masks could be lower quality in tissue regions with higher overall AT8 stain burden. To assess this, we analyzed whether the model performed differently in images with higher AT8 stain burden, using semi-quantitative scores and quantified diaminobenzidine (DAB) signals as correlative variables. We found the model exhibited consistent performance (mIOU) across images despite varying AT8 staining intensities (R^2^ = 0.109, Supp. Fig. 5c) and semi-quantitative grades (ρ = 0.026; Supp. Fig. 5c). Furthermore, the model effectively differentiated slide-level semi-quantitative grades assigned by an expert, outperforming grades assigned using DAB signal alone, particularly between the Moderate and Severe CERAD-like categories (Supp. Fig. 5b).

### Models identify NFT objects down to the pixel on one GPU an order of magnitude faster than humans can roughly point-annotate them

The conversion from the .czi file type to the .zarr file type averaged 7.7 minutes per WSI and required enough random-access memory (RAM) to load the entire uncompressed WSI (50.2 ± 21.4 GB). A user with RAM limits could use the aicspylibczi library instead to load regions of .czi files into memory rather than the entire image and build the Zarr files in successive chunks.

Generating a WSI-level segmentation takes approximately 32 minutes per WSI per GPU on a dataset with WSIs that average roughly 160,000 by 220,000px. This was 2.5x faster than reported by Wurts *et al.* 2020, who segmented WSIs of size 120,000 x 120,000 pixels at 20 minutes using four Volta V100 GPUs, which are comparable to the NVIDIA RTX 3090s we used. Further reduction in time could be achieved by generating a tissue mask via a lower-resolution image and running inference solely on the masked region. Generating WSI segmentations for the entire cohort (48 WSIs) took 12.8 hours across two NVIDIA RTX 3090s. The object detection model’s inference speed was approximately 60% faster, taking about 20 minutes per WSI per GPU; the entire cohort of 48 WSIs took 8.1 hours with 2 NVIDIA RTX 3090s.

To calculate the NFTDetector score, we loaded each segmented WSI at 1/64 of the original resolution and post-processed it through the blob and tissue detection pipelines (Methods). On average, this took 4 minutes and 1.5 minutes per WSI, respectively. The YOLOv8 model’s WSI-level score required a post-processing non-maximum suppression procedure to eliminate overlapping bounding boxes, averaging 5.2 seconds per WSI and totaling 4 minutes across the dataset.

The version-1 annotation files the trainee (KN) generated did not have timestamps per point annotation. However, through personal communications, we determined that the average ROI that contained NFTs took approximately 30 minutes to annotate. We estimate that generating the initial point annotations for this dataset took 33 hours (n = 1476 NFT annotations, or 44.7 annotations/hour). For the version-2 reannotation study, we directly parsed the image metadata from SuperAnnotate to assess the time it took for our expert annotator to add new point annotations to the dataset. Reannotating the entire dataset, which consisted of 975 “super-tiles,” took 82 minutes (11.9 super-tiles/minute). This reannotation process, which ultimately rescued 280 new annotations (or 204.9 annotations/hour), was completed in one sitting.

As the WSIs in the study had a mean of 1028 ± 957 detected NFTs, the model’s 1927.5 annotations/hour rate can automatically process entire slides at a ratio of 9.4x-43x faster than human annotators per single GPU.

### Slide-level neuropathologic burden scores from the NFT model correlated with separate expert semiquantitative assessments

While pixel-level identification of NFTs enables deep neuropathological phenotyping, many studies rely instead on WSI-level CERAD-like semiquantitative assessments of neuropathology burden. To calculate an automated per-slide single score for NFTs, we generated area-normalized scores from WSI-wide segmentations by counting the number of detected NFTs and normalizing the count by the tissue area (Fig. 5a). We then compared these calculated scores versus previously expert-assigned semi-quantitative scores and subjected them to statistical analysis using Spearman’s rho, ρ = 0.654 (Fig. 5b). The model accurately discriminated semi-quantitative categories, particularly in distinguishing Mild versus Moderate categories (Fig. 5b). As all cases had an intermediate/high AD neuropathologic diagnosis, few cases had none or very few NFTs; despite this, we qualitatively observed that the model discriminated between the None and Mild categories (i.e., very few detections in slides assigned None or Mild (Fig. 5c)).

For comparison, we also constructed an area-normalized WSI-score benchmark directly from the NFT point annotations provided by the novice using the same methodology as the model’s WSI-level scores. We used mixed effect models to characterize the difference in scoring between the model and annotator at each CERAD level while accounting appropriately for the strong similarity of samples from the same decedent. We found that both the model and the annotator scored the Mild-or-Less category significantly lower than the Moderate category, with a mean 38% lower for the novice (95% CI (0.15, 0.98)) and a somewhat greater drop for the model (24% vs. 38%, but not significantly different from the novice). Neither model nor novice showed statistically significant differences between samples from the Frequent and Moderate CERAD categories after accounting for repeated measurements from decedents. The model and novice did not score significantly differently from each other within the Moderate or Mild-or-Less categories; the estimated scores for the model were 63% higher in the Frequent category than for the novice, a near-significant difference (p = 0.061, 95% CI 2% less to 2.7-fold greater.)

Lastly, we performed a correlation analysis between demographic features, semi-quantitative scores, and WSI-level predictions. Categorical differences were not significant, except for the relationship between maximal educational attainment and whether the individual identified as Hispanic (p < 0.01, Supp. Fig. 10a). We observed strong positive correlations between the annotator’s tissue area-normalized score and both models’ tissue area-normalized scores per WSI (ρ_CNNv2_ = 0.704, ρ_YOLOv8_ = 0.773) and notable negative correlations between both models’ area-normalized scores and the age at death (ρ_CNNv2_= -0.375, ρ_YOLOv8_ = -0.405). We also observed a notable positive correlation between the annotator’s tissue area-normalized score and years of education but did not see the same results when comparing against either model’s scores (Supp. Fig. 10b).

## Discussion

We present a scalable deep-learning pipeline to detect the location and precise boundaries of mature neurofibrillary tangles (NFTs) in immunohistochemically stained whole slide images despite only using rapid single-point human annotations as training data. The models performed comparably to expert semi-quantitative slide-level scoring despite not being explicitly trained at this task and equivalently to the trainee annotations it learned on (Fig. 3b). The models also outperform both the trainee and the expert in NFT detection speed with pixel-level specificity and visually include only tangle-like objects in their predictions (i.e., other tau lesions such as tufted astrocytes, astrocytic plaques, or neuritic plaques are not predicted as mature tangles). Further, they demonstrate similar performance even as distinguishing pathologies from background signal becomes more difficult; more specifically, model performance remains strong as the AT8 burden in the images increases. We achieve this performance with a framework that is easily extended to other datasets and enables rapid integration and conversion of point annotations into either ground truth masks or bounding boxes. This pipeline exhibits stability in a cohort sampled across three different ADRCs and is likely to perform similarly on slides sourced from other institutions. Additionally, we publish the data, annotations, model weights, and easily extensible code along with the study if others wish to fine-tune the model or modify the pipeline to suit their dataset.

Studies using deep learning methods to detect NFTs predominantly omit segmentation models due to the massive annotator investment required to collect manual ground truth training masks at the pixel level[9,10,48] despite the reported research advantages of object- and pixel-based NFT counts over typical IHC positive pixel count methods[49]. Indeed, segmentation models unlock deeper neuropathological phenotyping than the standard pathological analyses that compress pathological information into a single category or semi-quantitative grade — examples include nuclei and tissue segmentation as tools to improve cancer grading schemes such as Gleason and C-Path scores[50–52]. Object detection models address some concerns, but the predicted bounding boxes they output do not directly capture morphological patterns in the detected NFTs. Although substantial morphological differences are well-documented between pretangles and mature tangles, NFTs may have undiscovered morphological nuances, particularly those characterized by distinct fibril structures, which are only unveiled through cryo-EM imaging[1,53]. Our study releases open-source NFT segmentation models trained solely from point annotations, requiring less annotator investment, delivering comparable object detection performance to published models, and exhibiting strong correlation with expert-assigned whole-slide image semi-quantitative grades[6,8,10,54,55].

Unlike other studies, we narrowed the focus to cells containing a nucleolus and mature tangles instead of annotating pre-tangles, ghost tangles, or cells that may appear neuronal but for which a nucleolus was not apparent. This likely decreased numerical model performance, as the decision boundary between mature tangles and other tau tangle categories is more complex than deciding between a tangle phenotype versus other objects or backgrounds in the slide. In addition, all annotations in this study are point annotations curated by a single trainee, then refined in an iteration by a single expert. By contrast, Signaevsky *et al*. perform NFT segmentation but report object-level F1 in their test set at 0.81. However, unlike our mature tangle annotation criteria, they include NFTs of various morphologies as valid annotations. The dataset included 22 cases of tauopathy sampled from the hippocampal formation and dorsolateral prefrontal cortex stained with AT8; ground truth mask annotations were manually generated across 178 ROIs by three expert neuropathologists (n_train_= 14 WSIs). In Vizcarra *et al.* 2023, a study involving various brain regions and multiple annotators (n_expert_ = 5, n_novice_= 3), the best object-detection macro F1 performance in the temporal region was 0.44 ± 0.06. This average considered pre-NFT and iNFT performance (F1_pre-tangle_ = 0.20, F1_iNFT_ = 0.71). Since we did not annotate and subcategorize pre-NFTs, they may be lumped into the model’s predictions for mature tangles; this may numerically penalize apparent model performance. Also, Vizcarra *et al.* note that there are generally more pre-NFTs predicted in the UC Davis cohort than iNFTs compared to Emory. In contrast, in their cohort, the opposite is true (albeit the model performed poorly in the UC Davis cohort overall). They proposed site-specific differences, such as differing tau antibodies. This lends evidence that our model suffers from the lack of pre-NFT labels. Their dataset, curated by five experts and three novices, underwent eight iterations of manual adjustment, removal, and addition of bounding boxes, whereas our study underwent a single iteration, collecting additional point annotations only.

Moving to object detection, Ramaswamy *et al*. 2023 report an amyloid plaque object detection average precision (AP) averaged over IOUs thresholds from 0.5 to 0.95 (AP@50:95) of 42.2. Wong *et al.* 2023 report 0.64 (model) vs. 0.64 (cohort) average precision (AP) for cored plaques and 0.75 vs. 0.51 AP for CAAs at a 0.5 IOU threshold (AP@50). We report an object detection AP@50:95 of 32.2 and AP@50 of 0.49 for mature NFTs programmatically bootstrapped from point annotations. In a third case, Koga *et al.* 2022 performed tau neuronal inclusion object detection in 2,522 CP13-stained images from 10 cases each of AD, PSP, or CBD across a range of brain regions; they achieved an average precision of 0.827. Koga *et al.* appear to use a more inclusive definition of tau neuronal inclusion by inspecting a representative lesion[55,56]. If we were to apply a similarly inclusive definition of NFT and hypothesize that 50% of our false positives were pre-NFTs or mature tangles outside of our annotation criteria, it would have a marked impact on the F1 score (from 0.58 to 0.74), but this is speculation.

We noted several caveats, which more typically arose from data than model architecture limitations. We could not assess whether the model can distinguish “None” versus “Mild” semi-quantitative categories due to the sample size (n=1 “None” test WSIs), although the model scored the “None” slide perfectly with an NFTDetector score of 0. Performance may vary by factors such as tissue staining variations due to overfixation or differences in scoring methodologies — semi-quantitative scoring analyzes the densest 1mm^2^ region of pathology, in contrast to the NFTDetector, which gives a WSI-level quantification. The standalone object detection model trained using bounding boxes derived from the ground truth segmentation masks likewise correlated with semi-quantitative grades to the same extent as the segmentation model. This suggested that both model strategies – segmentation and object detection – learned approximately equally well and possibly to their limit from the training data. Additionally, we focused on samples from individuals with pathologically defined AD, specifically from the temporal lobe, which may limit generalizability to other brain regions and disease presentations.

Notably, we release the first publicly available expert annotated dataset, trained model weights, and segmentation model codebase for NFT detection to support reproducibility and broad real-world use (https://github.com/keiserlab/tangle-tracer). These models can be used as-is, or the reader may wish to fine-tune or retrain them for institution- or goal-specific tasks. Instead of working directly with Python code, researchers and pathologists can use the platform without programming via its command line interface to convert new point annotations to segmentation ground truth masks (see the README.md in the code repository for details). We hope to integrate this pipeline and model with other pathology tools, such as the HistomicsUI[39] and active learning frameworks. Future studies may benefit from performing deeper morphological and correlative analyses between NFT features and clinical or genetic features using methods such as spatial transcriptomics.

## Conclusion

We introduce a robust and scalable deep learning approach to generate and detect ground truth masks for mature neurofibrillary tangles (NFTs). The segmentation model precisely detects and delineates mature tangles, enabling deeper morphological neuropathological analyses that may ultimately relate to subtle clinical and genetic factors. The model generates assessments comparable to expert semi-quantitative scoring when aggregated across an entire slide into a single score. Where speed is a concern, object detection models trained on bounding boxes calculated from the same data achieve similar but more rapid detection performance while giving up pixel-accurate NFT boundaries. Unlike other segmentation pipelines, the platform scales readily to new institutions because it “bootstraps” high-resolution NFT boundary training data automatically from simple and comparatively rapid human NFT single-point annotations. This tool reveals detailed maps of NFT distribution and morphology within tissue samples. We hope the open-source annotated dataset, trained model weights, and codebase we release here will facilitate diverse research directions, new uses, and collaborations in digital neuropathology.

## Supporting information

Additional File 3: Reannotation Protocol

## List of abbreviations

ML: machine learning
DL: deep learning
NFT: neurofibrillary tangle
WSI: whole slide image
Pre-NFT: pre-tangle stage of NFT
AD: Alzheimer’s Disease
PMI: Post-mortem interval aβ amyloid beta
DAB: diaminobenzidine
IHC: immuno-histochemical
NIA-AA: National Institute on Aging - Alzheimer’s Association
CERAD: Consortium to Establish a Registry for Alzheimer’s Disease
YOLO: You Only Look Once model
ADRC: Alzheimer’s Disease Research Center
ROI: region of interest
GPU: graphical processing unit
IOU: intersection over union
CLI: command line interface
TP/FP: true positive/false positive
ρ: Spearman’s rho

## Additional information

**Additional File 1 (.pdf):** Supplementary Material. Supplementary Figures and Tables.

**Additional File 2 (.xlsx):** Dataset 1. Spreadsheet containing Supplementary Tables 2-9. Additional File 1 contains descriptive captions for these Supplementary Tables.

**Additional File 3 (.pdf):** Reannotation protocol. Defines the reannotation procedure scope and provides instructions for using the SuperAnnotate platform.

## Declarations

### Ethics and approval and consent to participate

Ethical approval for this study was granted by the Institutional Review Boards at the home institutions of each ADRC, and written consent was obtained from individuals both during their lifetimes and posthumously. As the dataset originates from ADRCs, we consistently collected select data using standardized forms, ensuring data integrity and consistency across all cases.

### Consent for publication

Not applicable.

### Availability of data and materials

The images (rotated and cropped regions of interest and their segmentation masks) used to train and evaluate all machine learning models and data created and used in this project, including results, are available at https://www.ebi.ac.uk/biostudies/bioimages/studies/S-BIAD1165. The raw WSIs are available for download in Zarr format (must be uncompressed with imagecodecs’ Jpegxr codec; see repo below for example). The Python scripts, Jupyter notebooks, and additional code information are available at https://github.com/keiserlab/tangle-tracer. The README.md file contains detailed information on how to execute the pipeline and recreate results.

### Author contributions

Conceptualization: BND, MJK; Methodology: SG; Software: SG, LA; Validation: BND, LA, SG, SRRS, MJK; Formal analysis: SG, LA, LB, NS, MJK; Investigation: SG, BND, KN, LWJ, RR, AFT, CD; Resources: BND, MJK, CD, RR, AFT, LWJ; Data Curation: SG, BND, LM, KN; Writing - Original Draft: SG; Writing - Review & Editing: SG, BND, MJK; Visualization: SG, LA; Supervision: BND, MJK; Project administration: BND, MJK; Funding acquisition: BND, MJK.

## Acknowledgments

The authors thank the families and participants of the University of California Davis, University of California San Diego, and Columbia University Alzheimer’s Disease Research Centers (ADRC) for their generous donations, as well as ADRC staff and faculty for their contributions. This project was made possible by grant DAF2018-191905 (https://doi.org/10.37921/550142lkcjzw) from the Chan Zuckerberg Initiative DAF, an advised fund of the Silicon Valley Community Foundation (funder https://doi.org/10.13039/100014989) (M.J.K.) and grants from the National Institute on Aging (NIA) of the National Institutes of Health (NIH) under Award Numbers R01AG062517 (B.N.D.), P30AG072972 (C.D.), P30AG062429 (C.D.), P50AG008702 (A.F.T., Neuropathology Core), and P30AG066462 (A.F.T., Neuropathology Core). The views and opinions expressed in this article are those of the authors and do not necessarily reflect the official policy or position of any public health agency or the US government. The authors would also like to thank Mikio Tada and Irene Wang for their contributions throughout the project, as well as David Gutman and JC Vizcarra for sharing ideas and data to strengthen this study.

## Declaration of generative AI and AI-assisted technologies in the writing process

During the preparation of this work the author(s) used ChatGPT (https://chat.openai.com/, February 2024) to provide editing feedback for the manuscript and code (see Methods). After using this tool, the author(s) reviewed and edited the content as needed and take full responsibility for the content of the publication.

