## Additional File 1: Supplementary Figures and Tables for "Learning precise segmentation of neurofibrillary tangles from rapid manual point annotations"

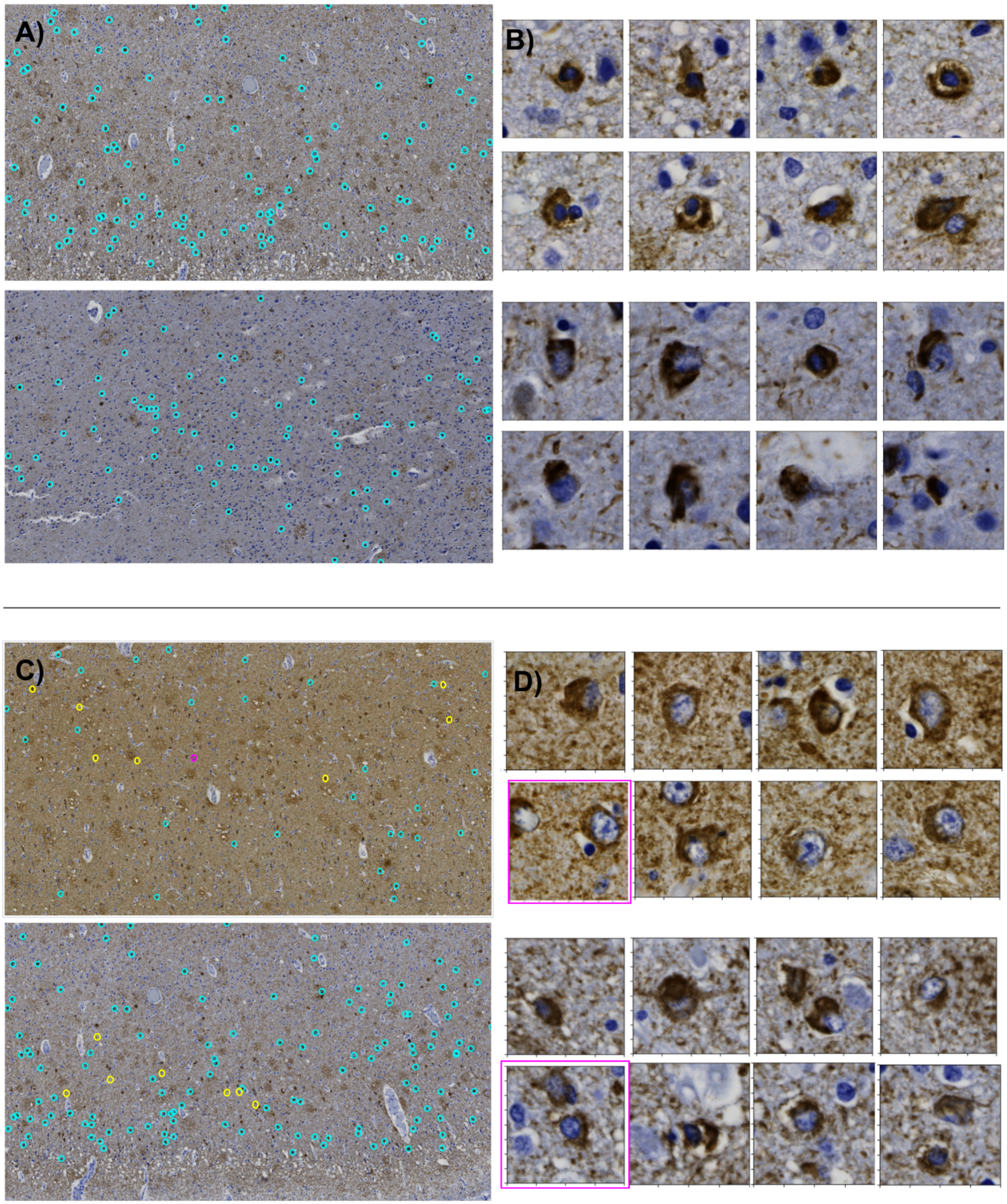
A)
B)
C)
D)

**Supp. Figure 1. Examples of mature tangle point annotations and NFT-like objects.** A) Two ROIs with cyan circles representing ground truth point annotations. B) Two groups of eight mature tangles derived from point annotations. C) Two ROIs where cyan circles are ground truth point annotations, yellow circles are tangle-like objects incorrectly labeled as NFTs, and pink circles are NFTs missed during reannotation but detected by the model. D) Two groups of eight mature NFTs where the boxes highlighted in pink correspond to the pink circles in the corresponding ROIs and the rest of the boxes correspond to yellow circles.

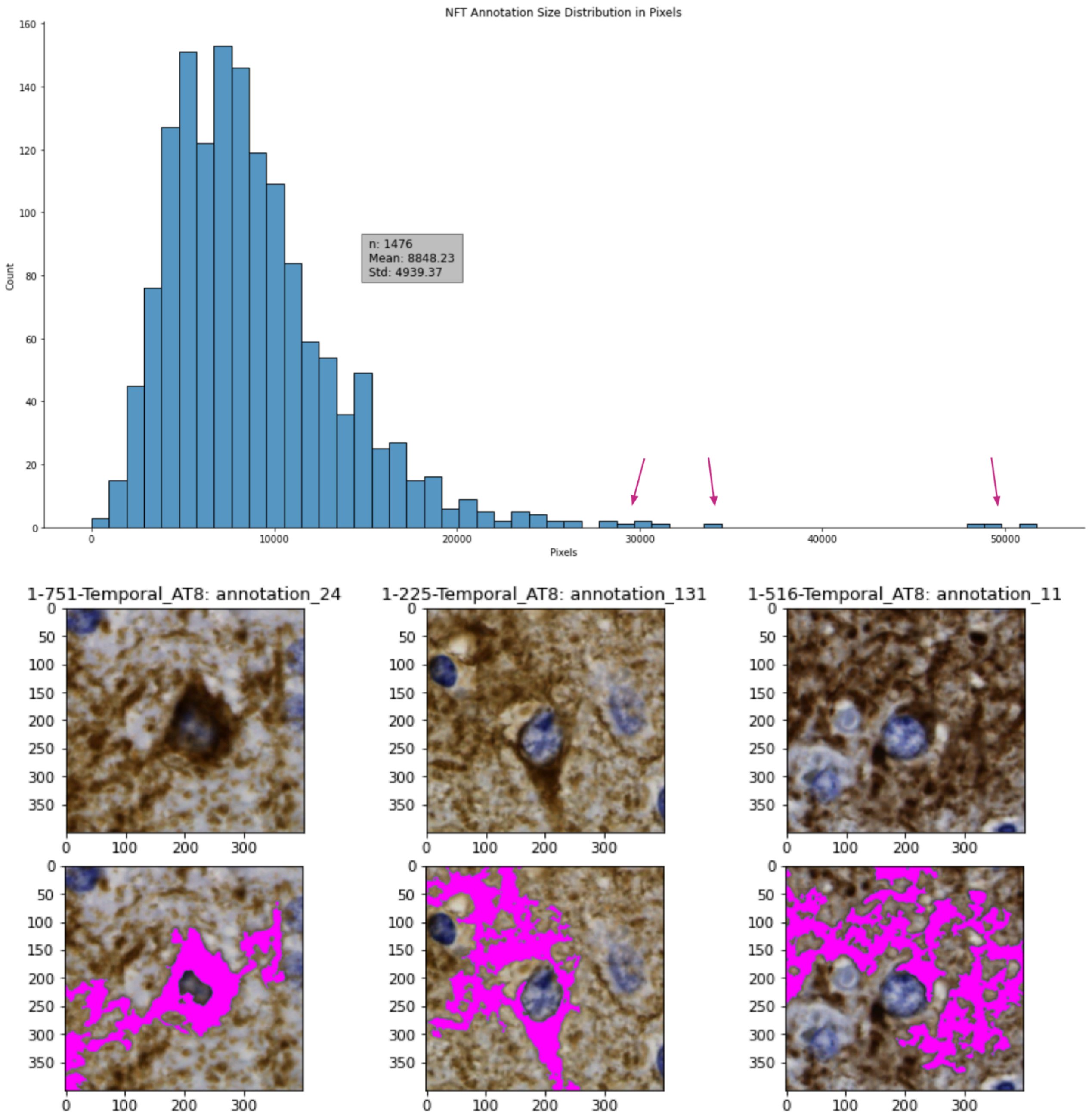


**Sup. Figure 2. Failure cases of pixel-to-mask (p2m) pipeline occur in predominantly high-background regions.** A) A histogram showing the distribution of NFT mask sizes in pixels. B) Examples of three NFTs across three different high-background WSIs where the point-to-mask pipeline did not generate masks within the boundary of the NFT. These failure cases were mostly observed in high-background ROIs.

A)

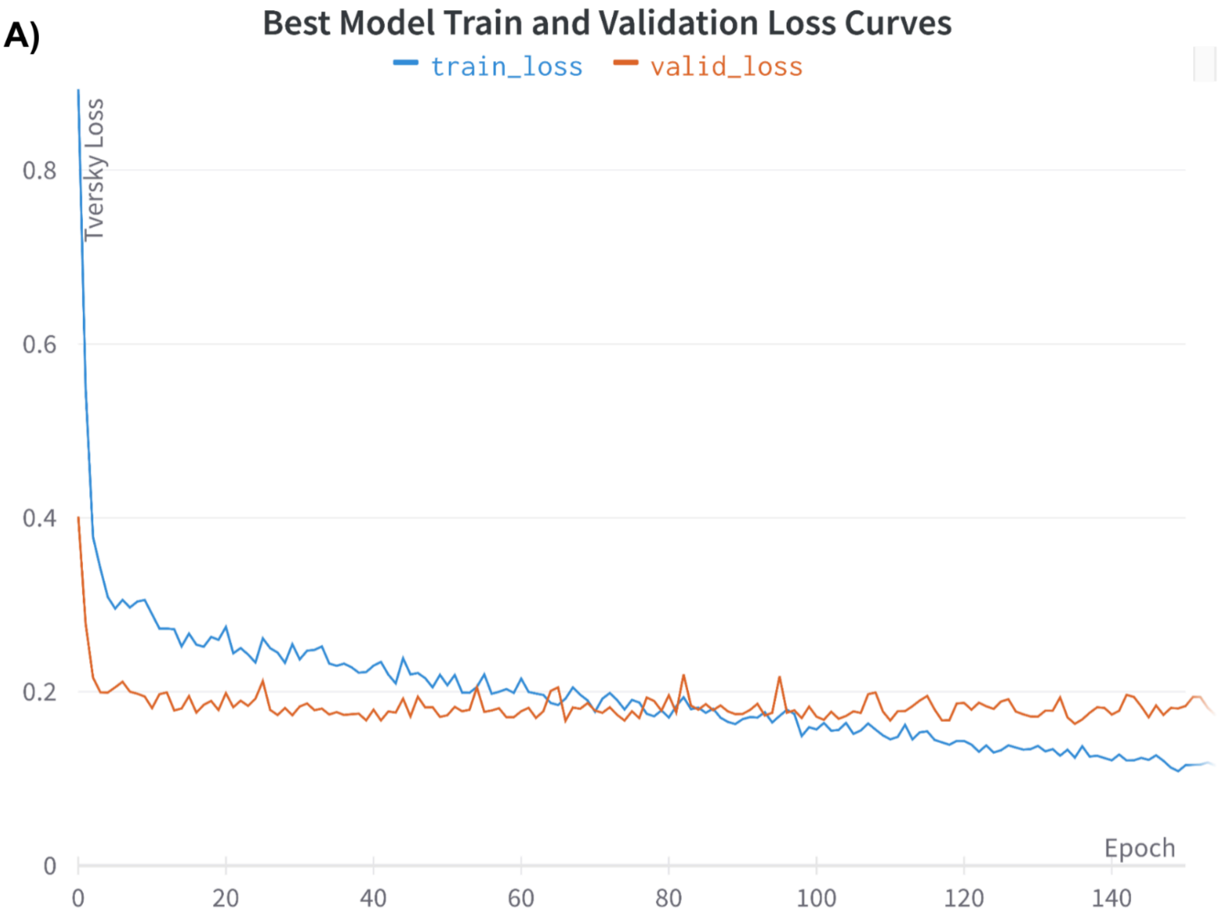
Best Model Train and Validation Loss Curves
train_loss
valid_loss
Tversky Loss
0.8
0.6
0.4
0.2
0
Epoch
0
20
40
60
80
100
120
140

B)

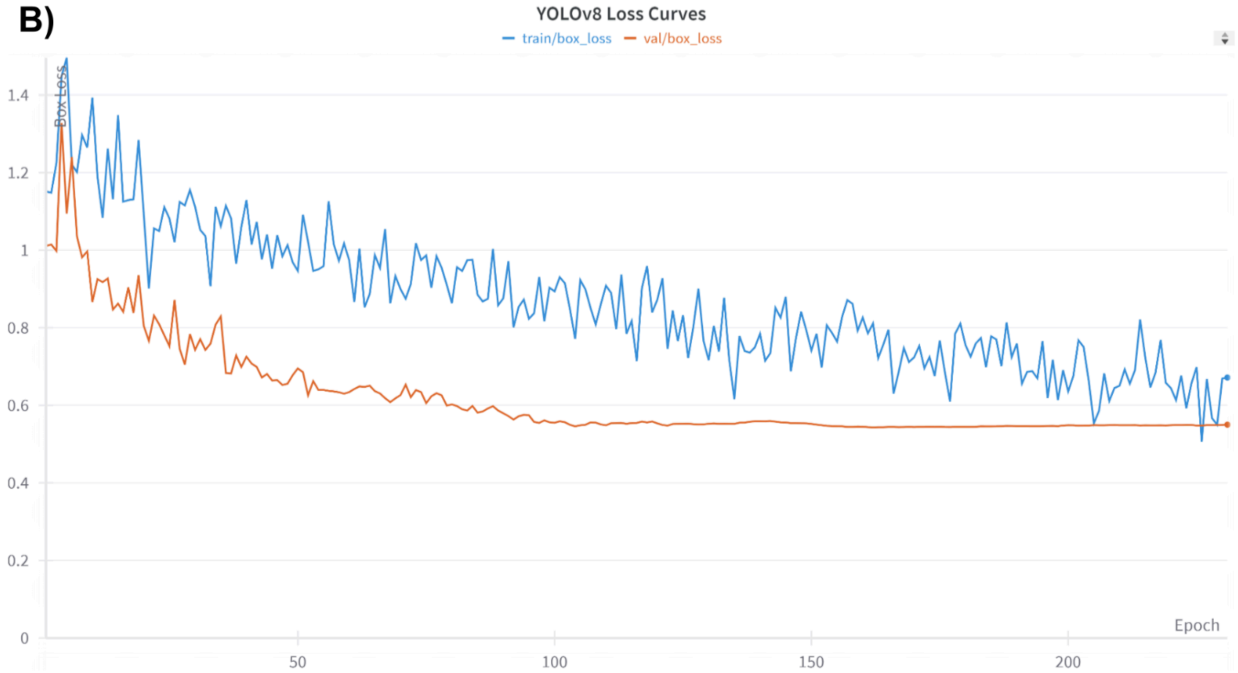
YOLOv8 Loss Curves
train/box_loss
val/box_loss
Box Loss
1.4
1.2
1
0.8
0.6
0.4
0.2
0
Epoch
50
100
150
200

**Supp. Figure 3. Loss curves for both models.** A) Validation and training loss curves (Tversky loss) for the segmentation model with the highest overall performance. B) Validation and training loss curves for the YOLOv8 object detection model with the highest overall performance. Displaying one of three logged losses, box loss.

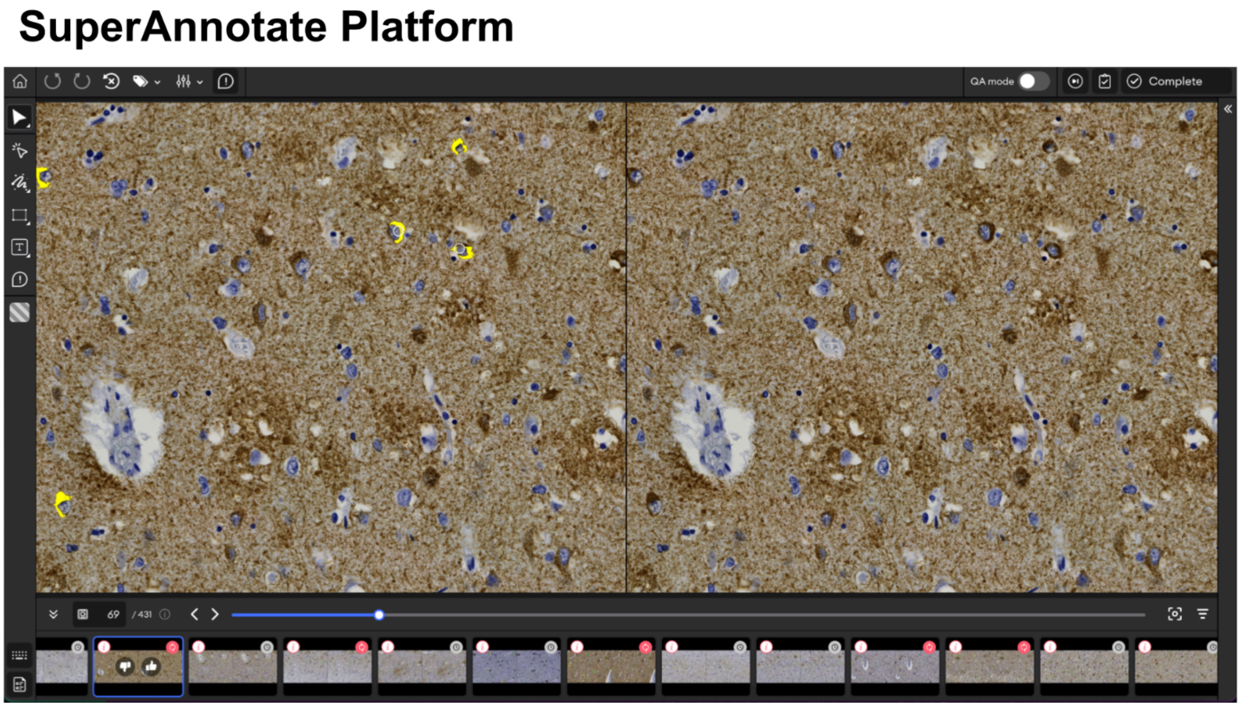


**Supp. Figure 4. Example of the SuperAnnotate platform.** The image on the right is the same as on the left but does not include the agreement map labels (yellow). The pathologist selected the points within the tile corresponding to NFTs that should be “rescued” and labeled as positive for the next iteration. No NFT annotations were removed using the platform.

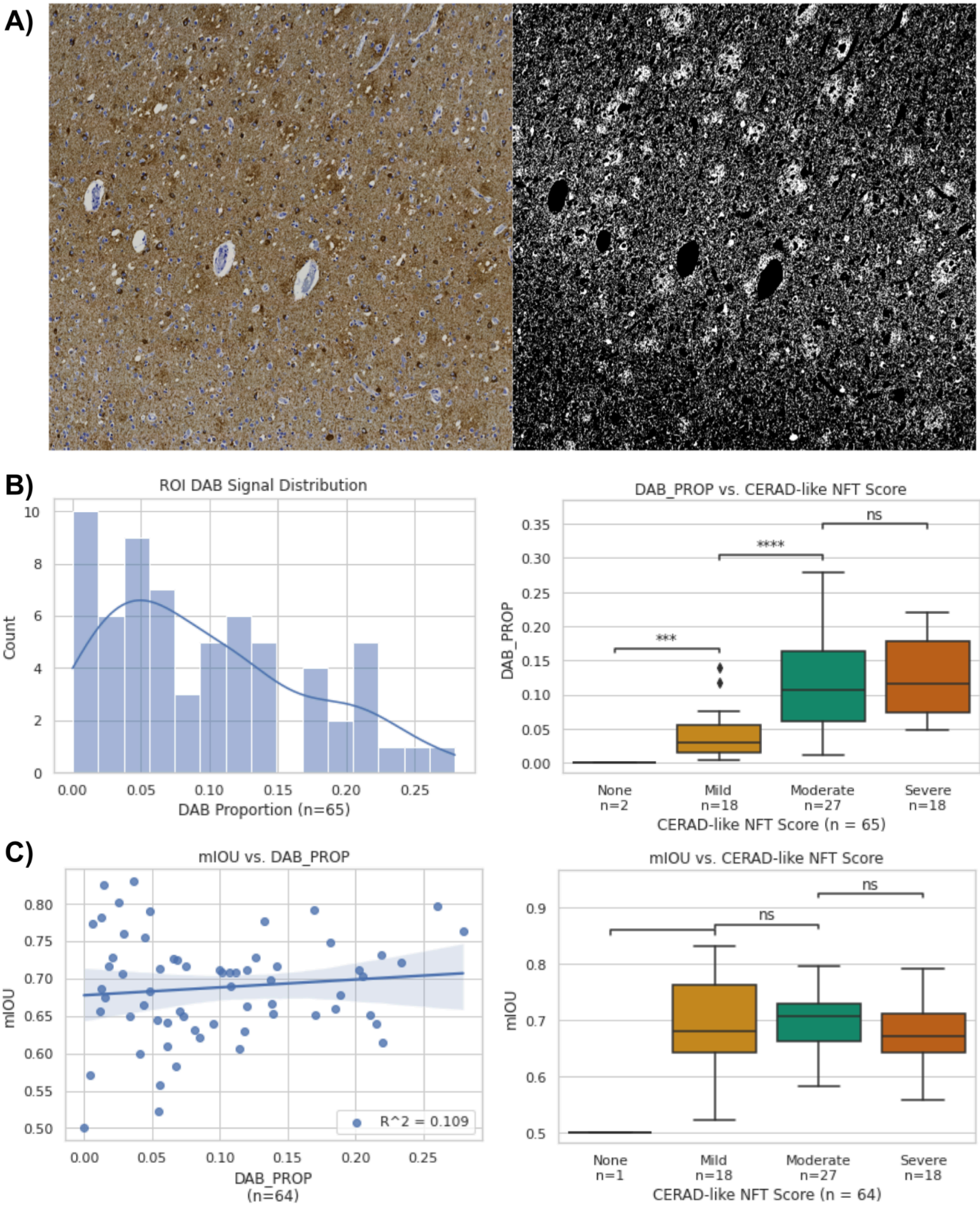


**Supp. Figure 5. Correlating DAB proportion and model performance.** A) An example ROI image crop (left) and the same image visualizing only the extracted DAB channel (right). B) Histogram (left) indicating the DAB signal proportion in each dataset ROI. Boxplot (right)

plotting the proportion of DAB signal against the semi-quantitative category independently assigned to the slide by a pathologist. We observed a linearly increasing relationship with a significant difference between the Mild and Moderate categories (Mann-Whitney U-Test). C) Scatterplot (left) with regression line plotting the model's mIOU for that ROI against the proportion of DAB in the ROI. We excluded the ROI with an undefined mIOU because it contained zero ground-truth NFTs. We find no significant correlation between these variables. Boxplot (right) plotting the mIOU of the model for a given ROI against its assigned semi-quantitative category. Again, we observe no significant correlation between these variables. ***: 0.0001 < p <= 0.001, ****: p < 0.0001 by Welch's t-test.

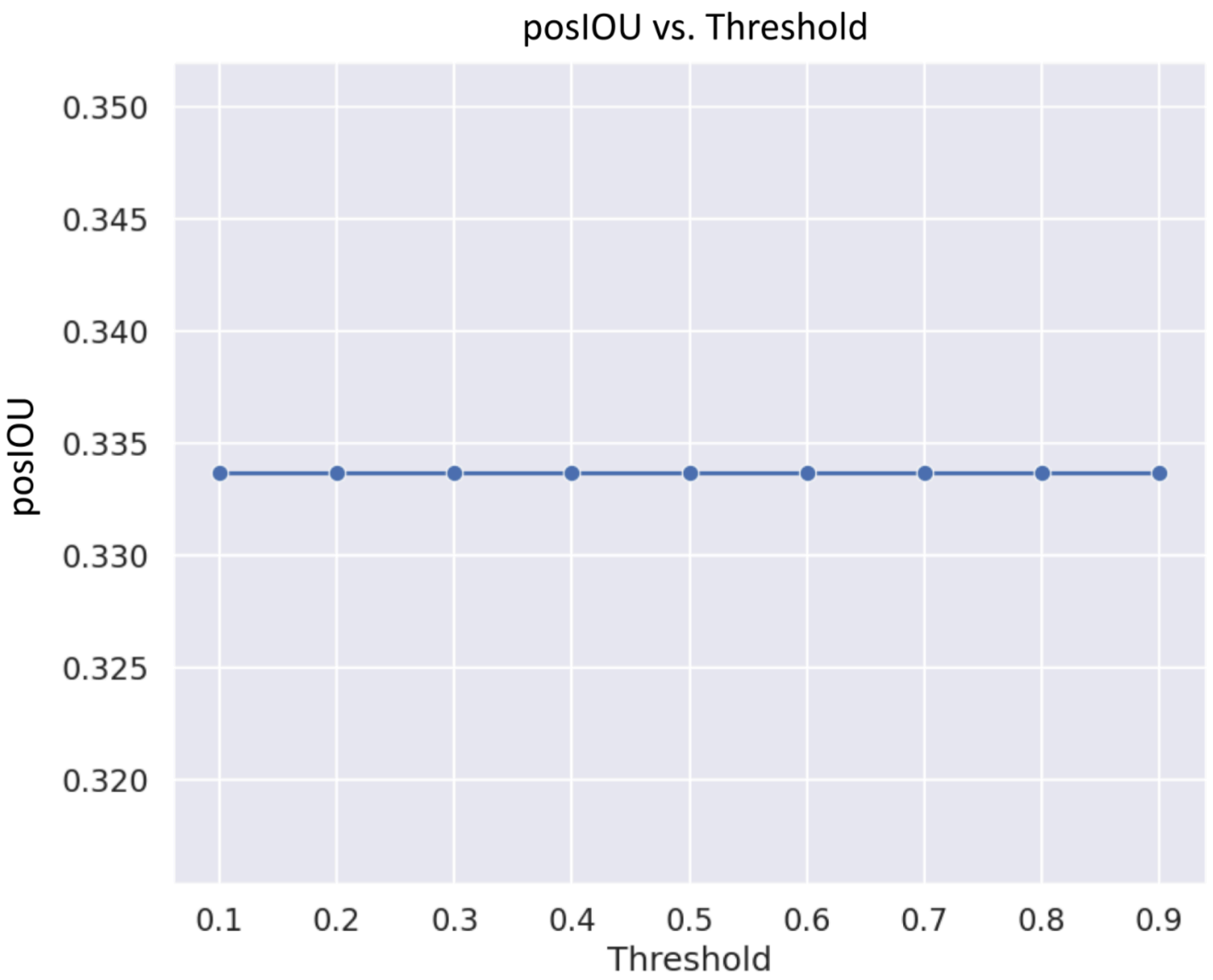


**Supp. Figure 6. Segmentation IOU vs Threshold.** Plot showing how positive IOU (and, therefore, mIOU) remained stagnant across IOU thresholds. Varying the prediction threshold when assessing ground truth and prediction overlap for segmentation did not affect performance.

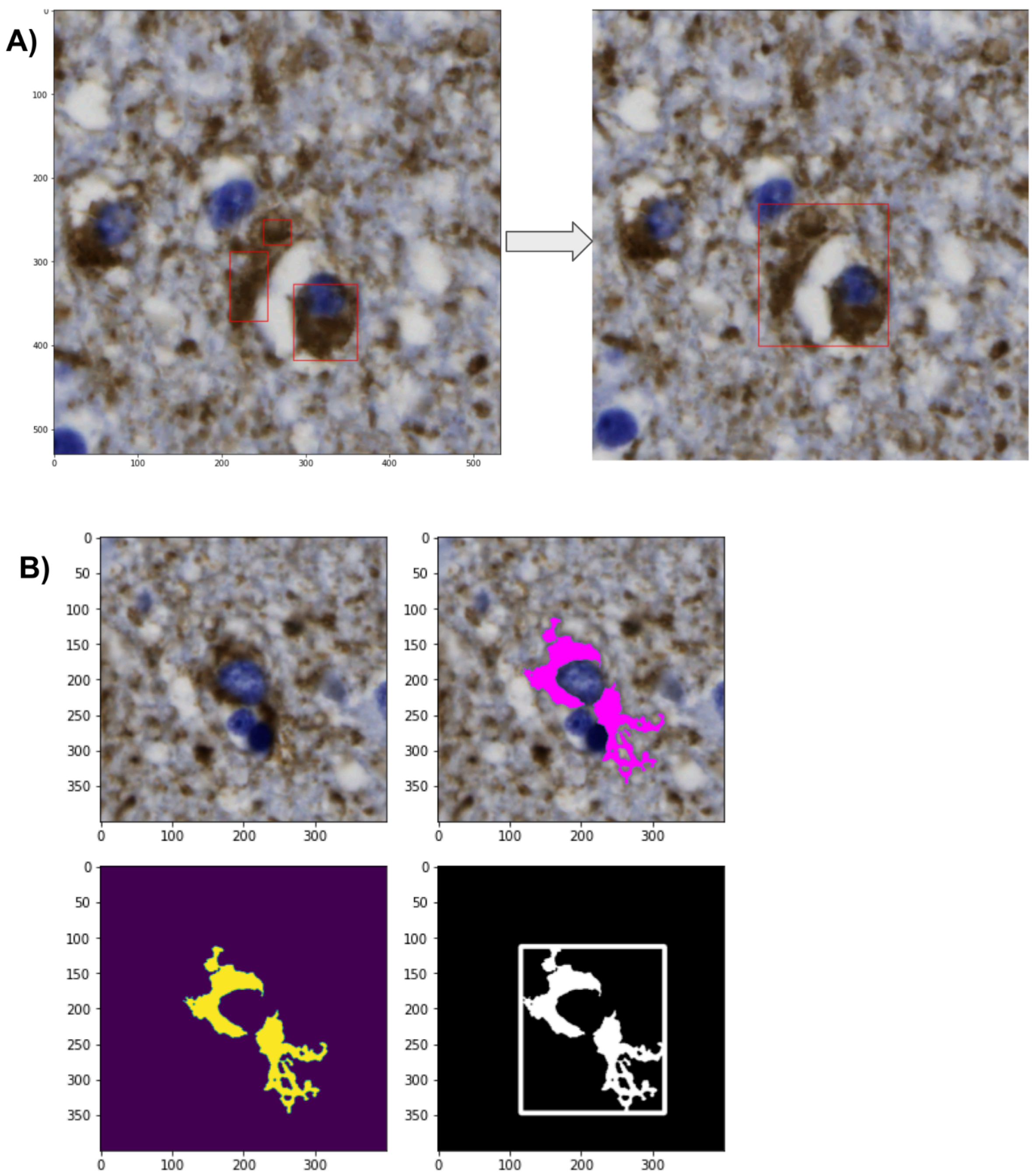


**Supp. Figure 7. Examples of converting segmentations to bounding boxes.** A) We merge bounding boxes encompassing different areas of the same NFT into one singular bounding box via a custom merging algorithm. B) One bounding box corresponds to one NFT mask in a 400x400 pixel crop. These are the same crops used to generate the masks for training the segmentation model.

A)

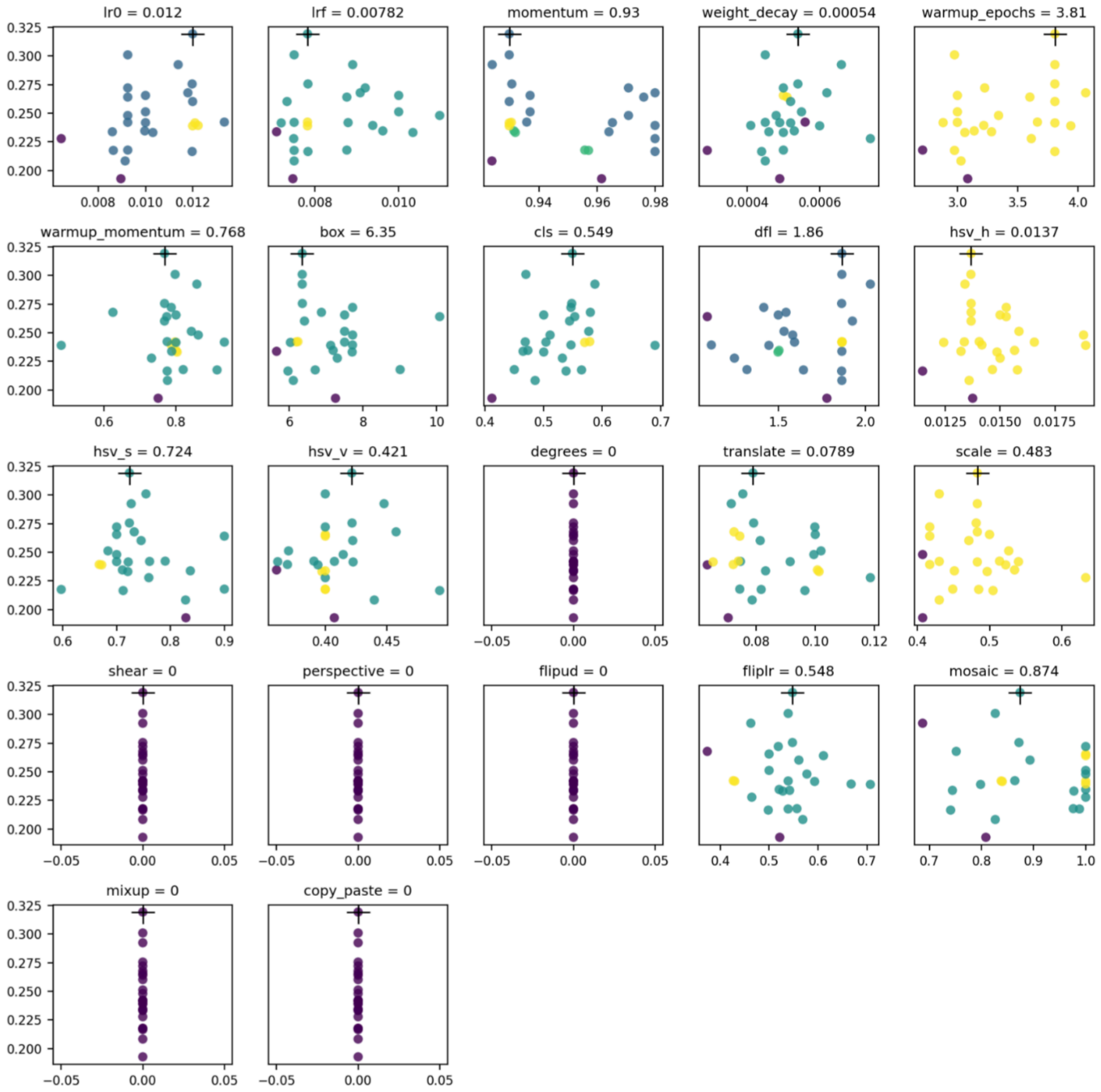
lr0 = 0.012
lrf = 0.00782
momentum = 0.93
weight_decay = 0.00054
warmup_epochs = 3.81
warmup_momentum = 0.768
box = 6.35
cls = 0.549
dfl = 1.86
hsv_h = 0.0137
hsv_s = 0.724
hsv_v = 0.421
degrees = 0
translate = 0.0789
scale = 0.483
shear = 0
perspective = 0
flipud = 0
fliplr = 0.548
mosaic = 0.874
mixup = 0
copy_paste = 0

B)

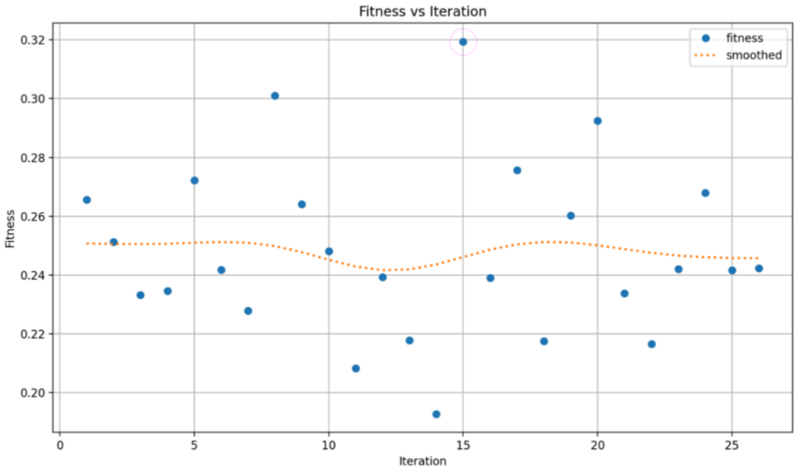
Fitness vs Iteration
fitness
smoothed
Fitness
Iteration

**Supp. Figure 8. Object Detection Hyperparameter Tuning.** A) Logged and plotted results of neural network model hyperparameter search. The variables included in the plot include a scatterplot with a point corresponding to a single iteration or model version with different parameters selected. B) Logged fitness of each model vs. iteration number. Fitness assessment is built into the Ultralytics library. The best model is circled in magenta.

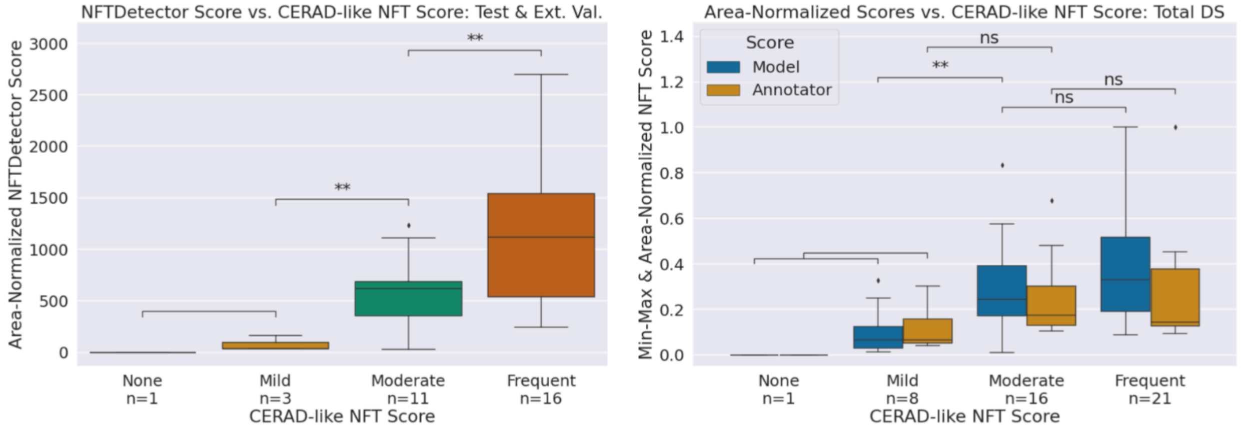


**Supp. Figure 9. Pre-reannotation performance metric box plots.** Two box plots for the model before the re-annotation experiment. One plot shows the area-normalized NFTDetector scores on the test set across increasing semi-quantitative scores (left-hand side), and the other shows a comparison of area-normalized NFTDetector scores on the entire dataset between the model and the annotator across increasing semi-quantitative scores (right-hand side). **: 0.001 < p <= 0.01 by Welch's t-test.

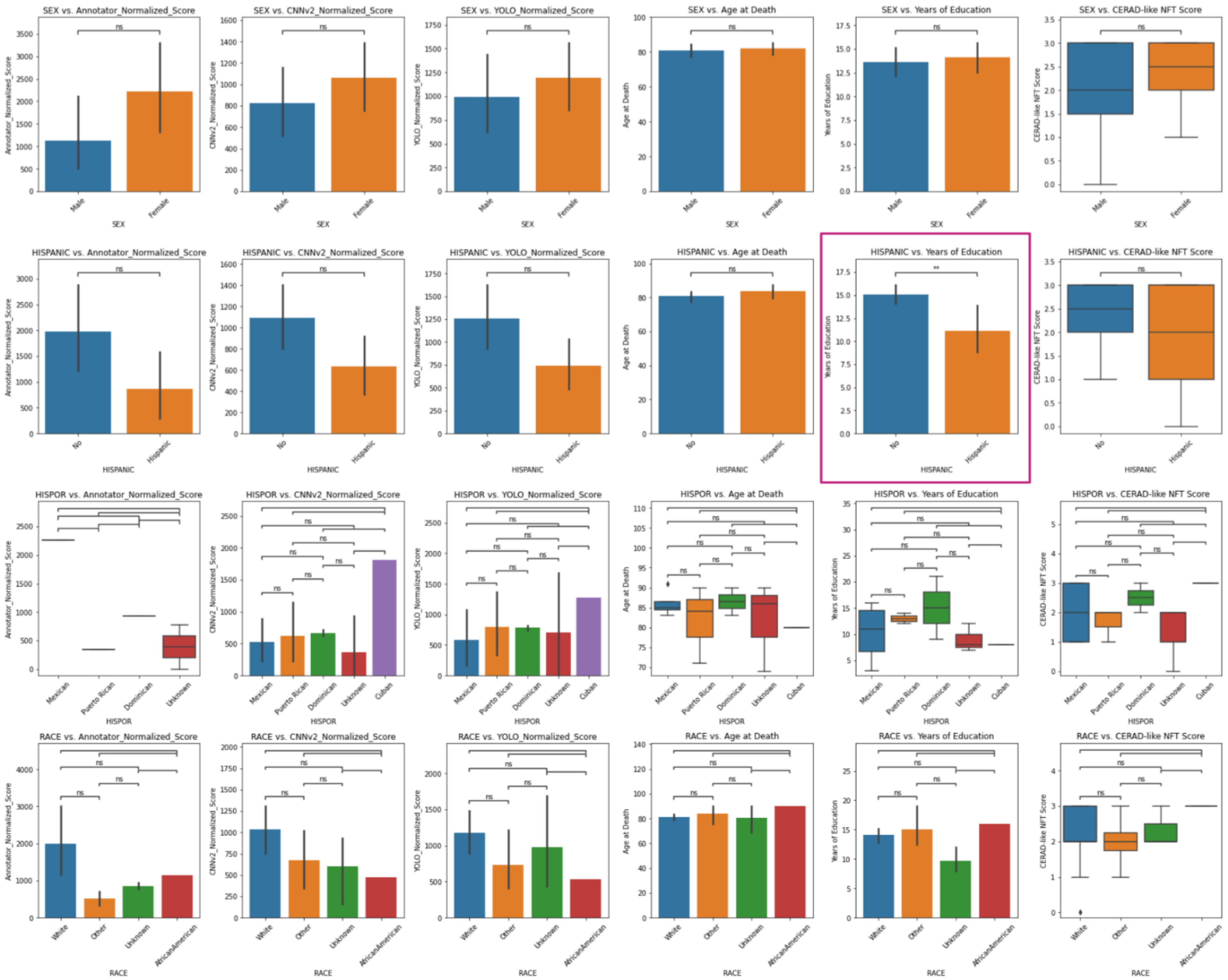


**Supp. Figure 10a. Correlative Analysis Between Demographic Features.** A collection of plots investigating the correlation of demographic features with slide-level information across all batch 1 WSIs ($n_{train/test} = 22$), including quantitative scores such as the models' scores (CNNv2_Normalized_Score, YOLO_Normalized_Score) and the number of annotations generated by the annotator (Annotator_Normalized_Score). Includes all categorical variables. **: $0.001 < p <= 0.01$ by Student's t-test.

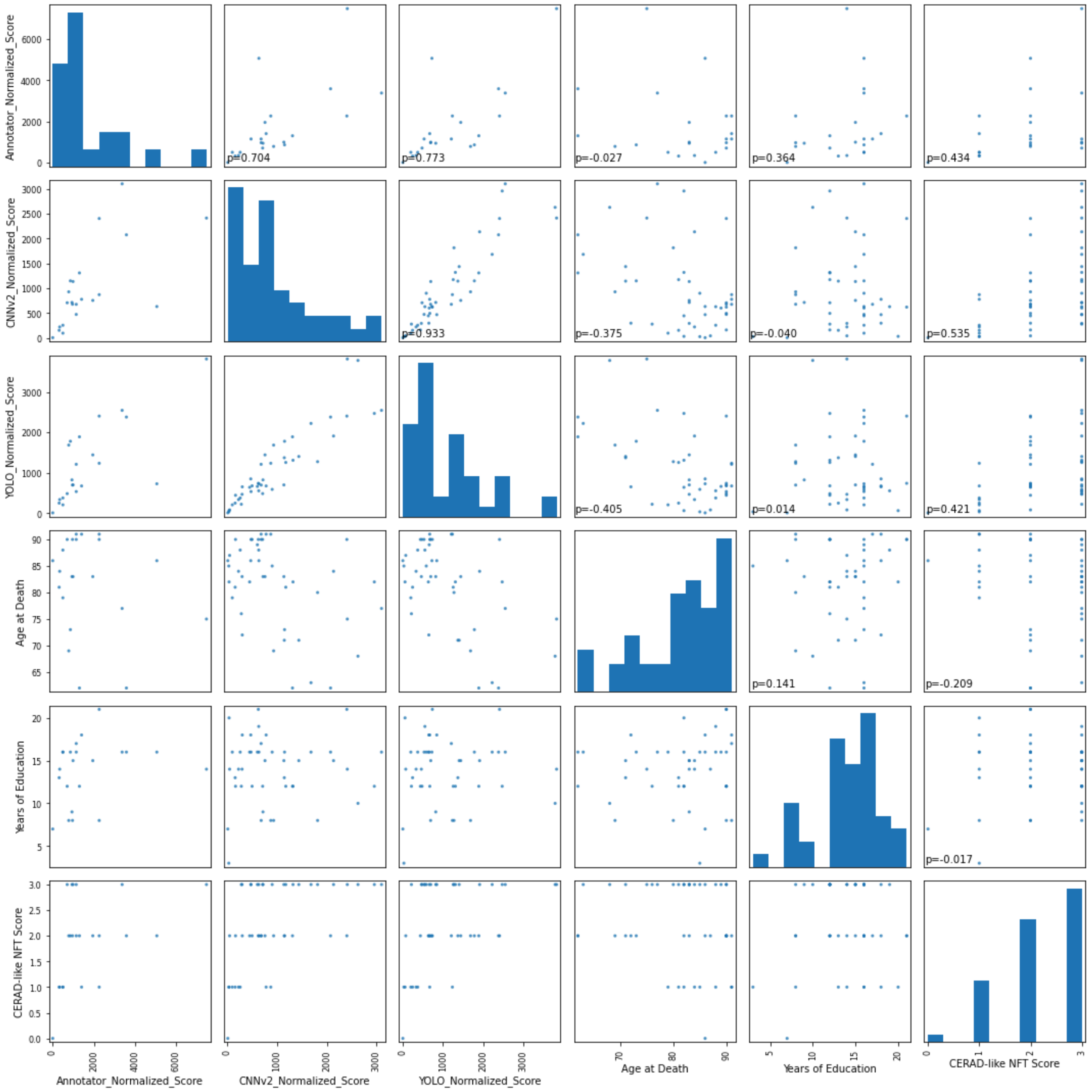


**Supp. Figure 10b. Correlative Analysis Between Demographic Features.** A collection of plots investigating the Pearson's rho correlation of demographic features against slide-level information across all batch 1 WSIs ($n_{train/test} = 22$), including quantitative scores such as the models' scores (CNNv2_Normalized_Score, YOLO_Normalized_Score) and the number of annotations generated by the annotator (Annotator_Normalized_Score). Includes all non-categorical variables.

### Supplementary Tables

| UNet model's object-detection performance | | | |
|---|---|---|---|
| **Dataset** | **Precision** | **Recall** | **F1** |
| Train | 0.346 | 0.952 | 0.507 |
| Val | 0.303 | 0.906 | 0.454 |
| **Test** | **0.464** | **0.682** | **0.552** |

**Supp. Table 1. Segmentation model's object-level metrics across dataset split**

### Supplementary Tables 2-9 Available in Additional File 2 (xlsx)

**Supp. Table 2.** Demographic information at the WSI (decedent) level. Demographic information was not given for case "02-996-Temporal_AT8," which we consequently excluded from Table 1. Rows highlighted in red correspond to WSIs not used in the study but were provided in either batches 1 or 2, as noted in Supp. Tables 8 and 9, respectively.

**Supp. Table 3.** Performance metrics at the ROI level for the UNet model trained after reannotation (v2). DAB_PROP: Diaminobenzadine signal expressed as the proportion of pixels positive. Precision, Recall, and F1 reflect the prediction accuracy at the pixel level (e.g., the proportion of the positive pixels recalled). posIOU: Positive intersection over union at the pixel level. In this case, positive refers to the NFT class. mIOU: An average of the positive and negative (background) IOUs.

**Supp. Table 4.** Object-level performance metrics at the ROI level for the UNet model trained after reannotation (v2). Results are calculated using bounding boxes drawn around ground truth and prediction masks. Performance metrics as described in Supp. Table 3 above but excludes posIOU and mIOU.

**Supp. Table 5.** Object-level performance metrics at the ROI level for the YOLOv8 model trained after reannotation (v2). Results are calculated using bounding boxes drawn around ground truth and prediction masks. Performance metrics as described in Supp. Table 3 above.

**Supp. Table 6.** Object-level performance metrics at the ROI level for the UNet model trained pre-reannotation (v1). Results are calculated using bounding boxes drawn around ground truth and prediction masks. Performance metrics as described in Supp. Table 3 above.

**Supp. Table 7.** All WSI-level dataset assignments (Train/Test), semi-quantitative scores (CERAD-like Score), tissue areas in pixels (Tissue_Area), and performance metrics for the annotator and each model trained (UNet and YOLOv8). Sets of performance metrics are in the following pattern: {Evaluation Method}_{Score Type} in sets of 3 corresponding to the raw score/count, the tissue area-normalized score, and the min/max tissue area-normalized score. For example, the count of NFTs generated by the annotator per ROI (Annotator_NFT_count), the annotator's tissue area-normalized score (Annotator_Normalized_Score), and the annotator's min/max tissue area-normalized score (Annotator_Min/Max_Norm_Score). Note: The tissue area used to calculate the annotator's normalized scores is calculated using the width and height of each ROI, not by the tissue area listed in the sheet, as that value corresponds to an entire WSI's area, not the ROIs' areas.

**Supp. Table 8.** Metadata describing each WSI (and its corresponding ROIs denoted in the annotations), available for download in batch1.zip (see Data Availability section). The USED column clarifies which slides were used in the study, and NOTES explains why certain WSIs were excluded. The train/val/test column denotes dataset assignment, and the NFT_PRESENCE column denotes whether any NFTs are present in the slide according to a novice annotator. Finally, the SCENE column denotes whether there is only 1 scene (None value) or multiple (numerical value). The WSI containing multiple scenes in this study was excluded.

**Supp. Table 9.** Table listing each WSI in the external validation batch, available for download in batch2.zip see Data Availability section). The NOTES column explains when WSIs were excluded from the study.
