## Additional File 3: Reannotation Protocol for "Learning precise segmentation of neurofibrillary tangles from rapid manual point annotations"

**Re-annotation experiment guidelines:**

- Link to SuperAnnotate project folder
    - The project folder is called ‘super-tiles’, which includes sub-folders containing the super-tiles (described below).
- Our goal is to ‘rescue’/annotate as many NFTs currently corresponding to the **yellow** (false positive) masks on the images in SuperAnnotate.
    - Adding a point annotation means you think the object is an NFT and should be labeled as an NFT in addition to the current set of NFTs (green and red).
    - If you do not think an object is an NFT, you do not need to annotate it or add anything to it – we can just leave it blank/as it is.
- Ignore green (true positive) masks completely
- Ignore red (false negative) masks and unlabeled (true negative) regions of the images **unless** something immediately looks like it should be annotated as an NFT. Otherwise, we don’t need to spend time annotating these portions of the tiles; we are primarily interested in the NFTs/NFT-like objects under the **yellow** masks.
- Please only drop point annotations on the **left-hand** tile (showing the agreement map). The right-hand tile/right-hand side of the whole uploaded image, which does not contain any labels/masks, is just the blank input image for your reference.
- Please drop point annotations roughly on/as close to the center of the NFTs as possible.
- Please annotate **as many supertiles as you can**. The super-tiles are divided into different folders on SuperAnnotate based on the order of priority in which we want to annotate them. Please go through the folders in the following order:
    - 1. A_test_fp
    - 2. B_train_fp
- These folders contain super-tiles where there are no/very small false-positive masks. These are optional/low-priority for annotation.
    - 3. C_test_no_fp (optional)
    - 4. D_train_no_fp (optional)

**SuperAnnotate features/tips:**

- You can check all of the keyboard shortcuts by clicking control+k/ctrl+k or clicking on the icon shown in the green circle below. However, the **main/important features are listed** in this document below.

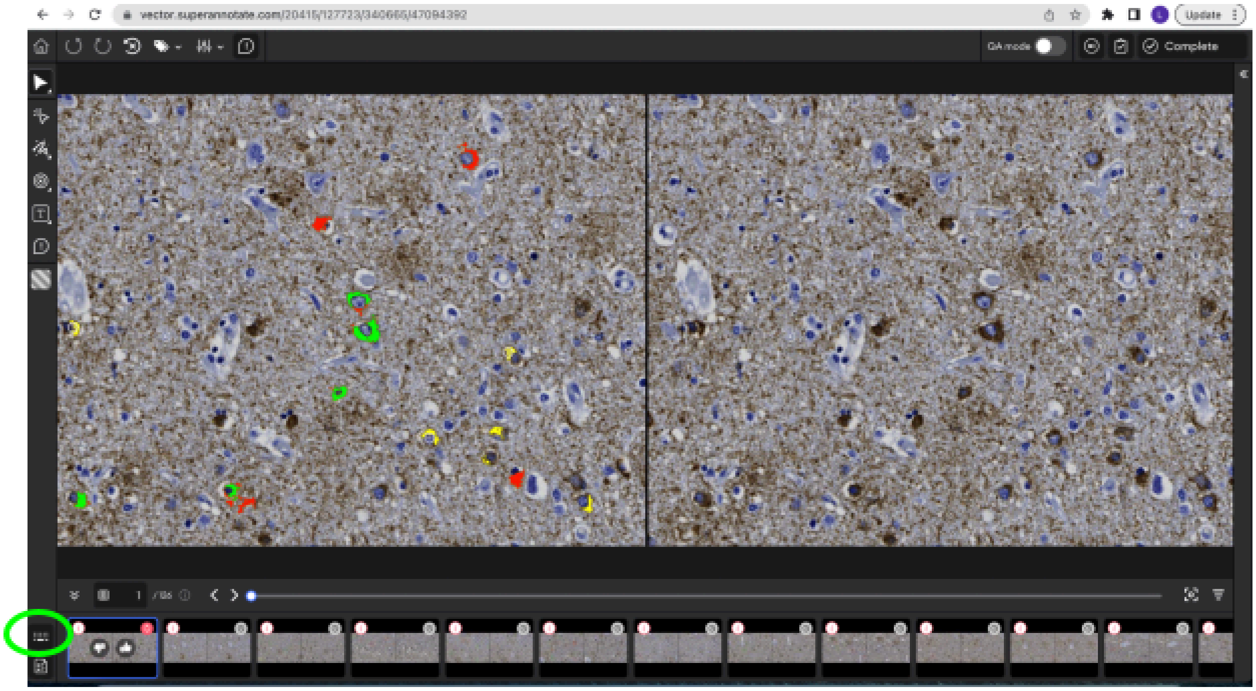


*Essentials*:

- **to make a point annotation:**
  - Press P on your keyboard and click on the part of the image where you want to drop the point annotation. The point annotation feature will toggle off automatically after you click on the image/drop the annotation.
  - You can also click on the bullseye-shaped icon on the left, shown below:

    - 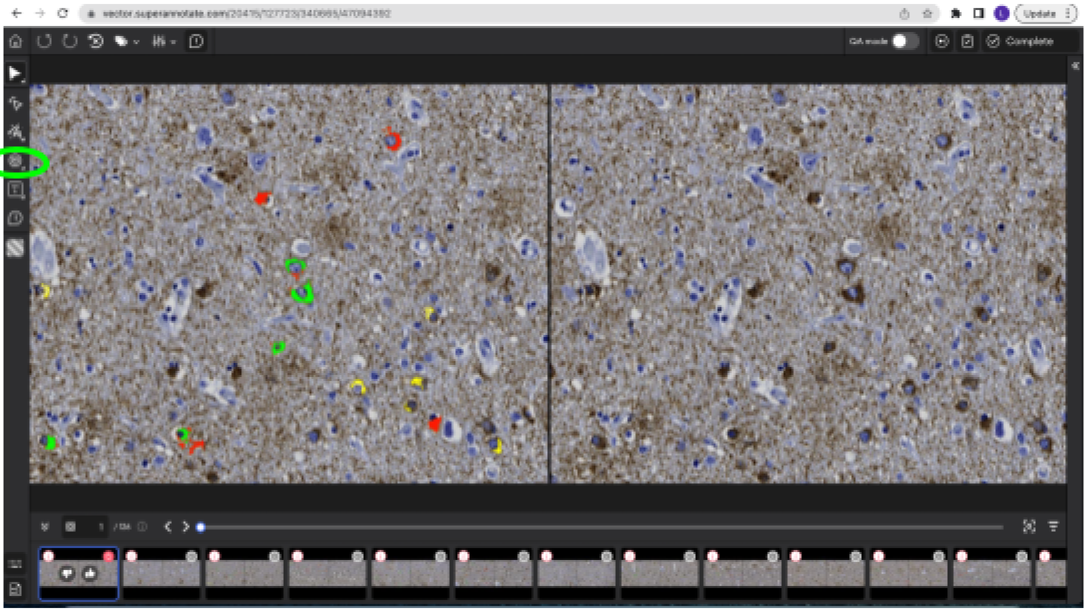


- to toggle the point annotation feature **off:**
    - Press V on your keyboard **or** press the mouse icon, shown below:
    - 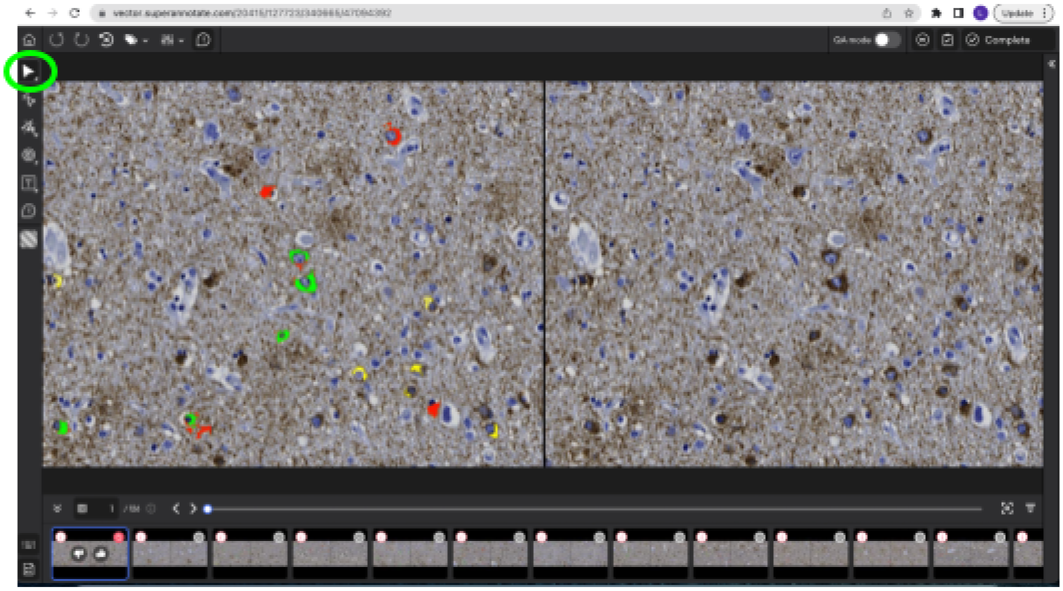


- **to zoom in:**
    - hover over the part of the image you want to zoom in to, and **scroll up** using your mouse/touchpad
- **to zoom out:**
    - **scroll down** using your mouse/touchpad
- **to go to the next image/previous image:**
    - Press the left/right arrows on your keyboard, or the left/right icons shown below:

    - 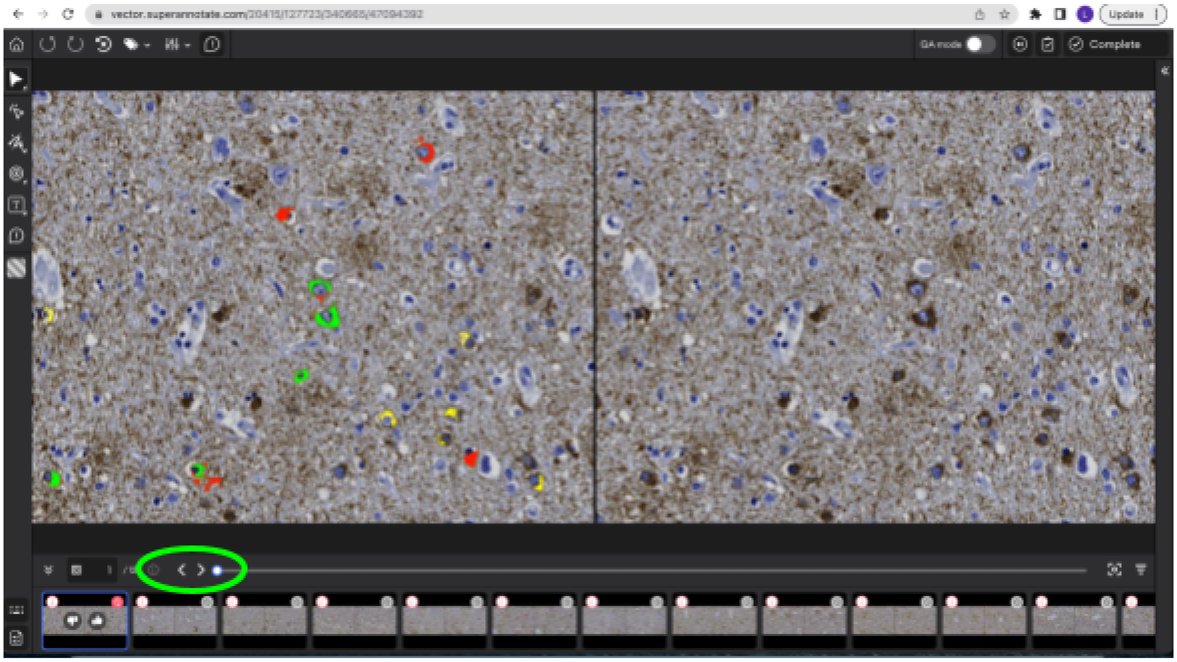


- **to save your annotations:**
    - ctrl+S (or control+S on Mac, **not** command)

*extra features, if they're helpful:*

- | | |
  |---|---|
  | = | Increase the brightness by 10% |
  | - | Decrease the brightness by 10% |
  | ] | Increase the opacity by 5% |
  | [ | Decrease the opacity by 5% |
  | Shift + + | Increase the contrast by 10% |
  | Shift + _ | Decrease the contrast by 10% |
